# The elasticity of individual protocadherin 15 molecules implicates cadherins as the gating springs for hearing

**DOI:** 10.1101/503029

**Authors:** Tobias F. Bartsch, Felicitas E. Hengel, Aaron Oswald, Gilman Dionne, Iris V. Chipendo, Simranjit S. Mangat, Muhammad El Shatanofy, Lawrence Shapiro, Ulrich Müller, A. J. Hudspeth

## Abstract

Hair cells, the sensory receptors of the inner ear, respond to mechanical forces originating from sounds and accelerations^1,2^. An essential feature of each hair cell is an array of filamentous tip links, consisting of the proteins protocadherin 15 (PCDH15) and cadherin 23 (CDH23)^3^, whose tension is thought to directly gate the cell’s transduction channels^4,5,6^. These links are considered far too stiff to represent the gating springs that convert hair-bundle displacement into forces capable of opening the channels^7,8^, and no mechanism has been suggested through which tip-link stiffness could be varied to accommodate hair cells of distinct frequency sensitivity in different receptor organs and animals. As a consequence, the gating spring’s identity and mechanism of operation remain central questions in sensory neuroscience. Using a high-precision optical trap, we show that an individual monomer of PCDH15 acts as an entropic spring that is much softer than its enthalpic stiffness alone would suggest^7,8^. This low stiffness implies that the protein is a significant part of the gating spring that controls a hair cell’s transduction channels. The tip link’s entropic nature then allows for stiffness control through modulation of its tension. We find that a PCDH15 molecule is unstable under tension and exhibits a rich variety of reversible unfolding events that are augmented when the Ca^2+^ concentration is reduced to physiological levels. Tip-link tension and Ca^2+^ concentration are therefore likely parameters through which nature tunes a gating spring’s mechanical properties.

---

Mechanically gated ion channels are ubiquitous. In addition to underlying our senses of hearing, balance, and touch, they are involved in the regulation of processes such as muscle extension, blood pressure, pulmonary inflation, and visceral distension. These channels are opened and closed through the action of gating springs, which are elastic elements that are tensioned by mechanical stimulation and in turn communicate stress to the molecular gates of the respective channels. Gating springs accordingly store mechanical energy and use it to regulate channels’ open probabilities. For bacterial mechanoreceptors, which respond to osmotic stress, the cellular membrane itself serves as a gating spring^9^. The ubiquitous Piezo channels of vertebrates extend three membrane-embedded arms that likely act as gating springs by flexing in response to membrane stretching^10,11^. Other mechanosensitive channels, such as NOMPC (TRPN1) in *Drosophila*, appear to be gated by the tension in elastic ankyrin domains^12^.

Gating springs were first posited for hair cells of the vertebrate inner ear, the sensors of the auditory and vestibular systems^4^. Each hair cell is surmounted by a hair bundle—a cluster of erect, actin-filled processes termed stereocilia—that is deflected by mechanical stimulation. However, the identity of the gating springs in these cells has remained controversial. A plausible candidate discovered soon after the gating-spring hypothesis was advanced is the tip link. Extending about 150 nm between the tip of each stereocilium and the side of its longest neighbor, the tip link is positioned to sense the shear between stereocilia when a hair bundle is deflected (Figure 1a,b)^5,13^. The tip link is a dimer of parallel dimers, comprising two PCDH15 molecules joined at their amino termini to a pair of CDH23 molecules through a “handshake” whose stability depends upon the presence of bound Ca^2+^ ions (Figure 1c)^14^. The mechanical properties of hair bundles imply a gating-spring stiffness^15,16^ on the order of 1 mN·m^-1^. However, electron-microscopic images suggest that the tip link is relatively rigid^17^ and crystallographic studies and molecular-dynamics simulations of the relevant cadherins support a stiffness fiftyfold as great as that measured^7^. It has therefore been posited that most of a gating spring’s elasticity resides at a tip link’s two attachments, rather than within the link itself. To clarify the identity of the hair cell’s gating spring, we have therefore examined the elastic properties of a tip-link protein.

**Fig. 1.**
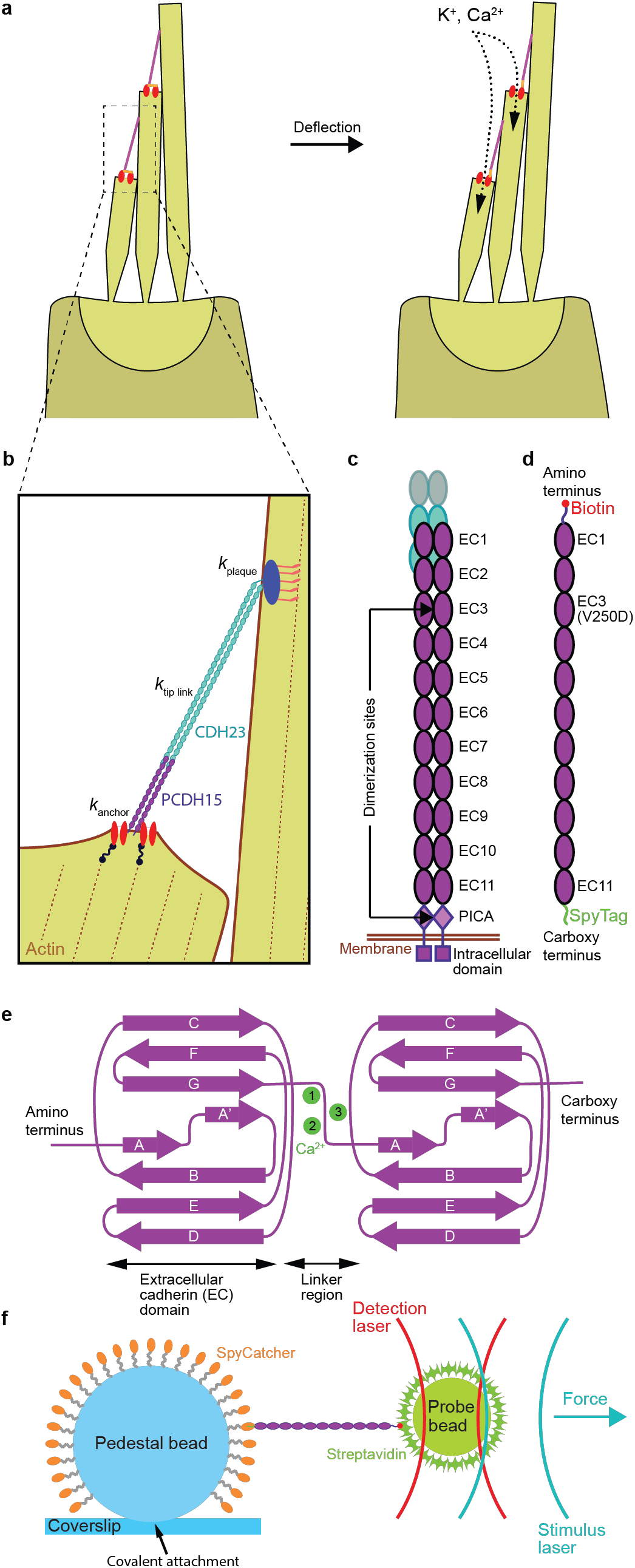
The role of tip-link proteins in transduction by hair cells. **a**, The hair bundle is a cluster of stiff, actin-filled protrusions called stereocilia that stands atop each hair cell in the inner ear. Each stereocilium is connected to its tallest adjacent neighbor through a proteinaceous filament called a tip link (pink), which is coupled at its base to mechanically gated ion channels (red). Deflection of a hair bundle increases the tension in the tip links, biasing the channels toward an open state that allows the influx of positively charged ions. **b**, The mechanical element that converts hair-bundle displacement into a force capable of opening the channels is called the gating spring. Its stiffness comprises those of the channel and its lower anchor (*k*_anchor_), the tip link proteins PCDH15 and CDH23 (*k*_tip link_), and the insertional plaque that anchors the link’s top end into the taller stereocilium (*k*_plaque_). **c**, The mechanical properties of the tip link emerge from its quaternary structure and from the characteristics of its constituent proteins. The lower third of the link consists of a dimer of PCDH15 molecules, each of which includes eleven extracellular cadherin (EC) domains. **d**, To measure the mechanical behavior of monomeric PCDH15, we tagged each end with a distinct molecular handle. We eliminated dimerization by a point mutation (V250D) in domain EC3 and by truncation of the PICA domain. **e**, The folding motifs of individual EC domains influence the mechanical properties of the full-length protein. Up to three calcium ions (green) can bind between domains. **f**, We probe the mechanics of a PCDH15 monomer by confining it through its handles between an immobile, 2 μm glass pedestal bead and a diffusive, 1 μm plastic probe bead. To acquire each force-extension relation, we measure the position of the probe bead with a detection laser while applying a force with a stimulus laser.

## Measurement of PCDH15’s mechanical characteristics

The mechanics of a protein can be tested by tethering it between two surfaces, applying a force that pulls the surfaces apart, and measuring the protein tether’s elongation^18^. To explore directly the stiffness of a tip link, we used a high-precision optical trap to determine the mechanical properties of individual molecules of PCDH15. The extracellular portion of PCDH15 consists of eleven extracellular cadherin domains (EC1-EC11) followed by the membrane-proximal PICA domain. PCDH15 forms a homodimer through interfaces in EC3 and in PICA (Figure 1c)^19,20^, hampering our study of the monomeric protein. We therefore introduced a dimerization-disrupting mutation (V250D)^19^ into EC3 and truncated the protein just before the PICA domain (Figure 1d).

Using a short, site-specific anchor at each of the protein’s ends, we placed PCDH15 between an immobile, 2 μm-diameter glass pedestal bead and a mobile 1 μm probe bead. Short, stiff anchors were necessary to avoid masking the elastic properties of PCDH15 by those of the anchors (Figure 1f). The probe bead was confined in a weak optical trap whose forward-scattered light we collected to determine the three-dimensional position of the bead with sub-nanometer spatial and 1 μs temporal resolution^21,22^. A second optical trap, displaced by a few hundred nanometers from the first, served to deliver a force stimulus. By increasing the trap’s stiffness, we could apply forces up to and exceeding 60 pN to the tethered protein (Figure 1f).

Cadherin domains manifest a stereotyped, immunoglobulin-like Greek key folding motif and are separated from one another by conserved linkers that each bind up to three Ca^2+^ ions in a canonical arrangement (Figure 1e)^23^. Ca^2+^ binding is thought to rigidify the linker regions and to stabilize the cadherin domains against force-induced unfolding that elongates the molecule^24^. The Ca^2+^ concentration of the endolymph, the fluid surrounding tip links, varies between different organs and species from tens to hundreds of micromolar^25,26^. Such concentrations are close to the Ca^2+^ dissociation constants for the various binding sites^7,27^, raising the intriguing possibility that nature adjusts the Ca^2+^ concentration to tune the mechanics of tip-link cadherins. We therefore explored the behavior of tensioned PCDH15 at three Ca^2+^ concentrations: 3 mM, to saturate all binding sites; 20 μM, a concentration that mimics the concentration of Ca^2+^ in the endolymph of the mammalian cochlea^25^; and zero.

Because structural changes in proteins are stochastic events that are driven by thermal forces, the rate at which external force is applied can dramatically change the mechanical response. If a protein is pulled too fast, thermal forces do not have sufficient time to cause barrier crossing in the protein’s energy landscape before very high external forces are reached. Consequently, the forces at which structural changes occur are artificially elevated at high loading rates^28^. Even in the absence of an acoustic stimulus, tip links experience a constant resting tension^29^ that varies with the frequency sensitivity of a hair cell, from 8 pN at 1 kHz to 50 pN at 4 kHz, with possibly even greater tensions at higher frequencies^16^. In each tip link two copies of PCDH15 act in parallel, and each copy assumes half of the tension of the entire link, 4 pN up to at least 25 pN. During normal hearing, a sound stimulus then superimposes an oscillation of only a few piconewtons on this resting tension. Our experiments therefore had to apply slowly changing forces to explore the influence of resting tension on the mechanics of PCDH15.

We applied force ramps to the single-molecule tethers at a loading rate of 130 pN·s^-1^ (unless noted otherwise) by linearly increasing the spring constant of the stimulus trap (Extended Data Fig. 1). This rate represented a compromise between slow force application and our desire to collect a statistically relevant number of extension-relaxation cycles for each molecule tested in a reasonable amount of time. For each cycle, we ramped the force up to 60 pN to cover the entire range of physiologically relevant tensions, then returned it at the same rate to a holding level of 2-4 pN. Depending on the Ca^2+^ concentration, we adjusted the holding level and duration to allow the protein to refold domains after many but not all cycles. The chosen loading rate likewise led to unfolding events in only a subset of cycles. With these parameter choices we were able to trap the protein in a given conformational state for several extension-relaxation cycles, allowing us to precisely characterize the mechanics of each state.

Figure 2a-c shows representative examples of individual force-extension relations for Ca^2+^ concentrations of 3 mM, 20 μM, and zero. Each curve features a hockey stick-like shape, as expected for the extension of a biopolymer in a heat bath^30^. As we quantify below, this functional shape indicates that entropic effects dominate PCDH15’s elastic response. Abrupt, stepwise extensions or “rips” in the force-extension relations correspond to structural changes of the protein under force. In contrast to typical single-molecule experiments, under our loading conditions PCDH15 never fully unfolded during the extension phase of the stimulus. We therefore frequently observed extensional structural changes even in the relaxation phase of the stimulus. For each Ca^2+^ concentration, a set of conformational changes leads to a modulated occupation of the force-extension state space, which we visualize by overlapping hundreds of extension-relaxation cycles for one representative molecule apiece (Figure 2d-f).

**Fig. 2.**
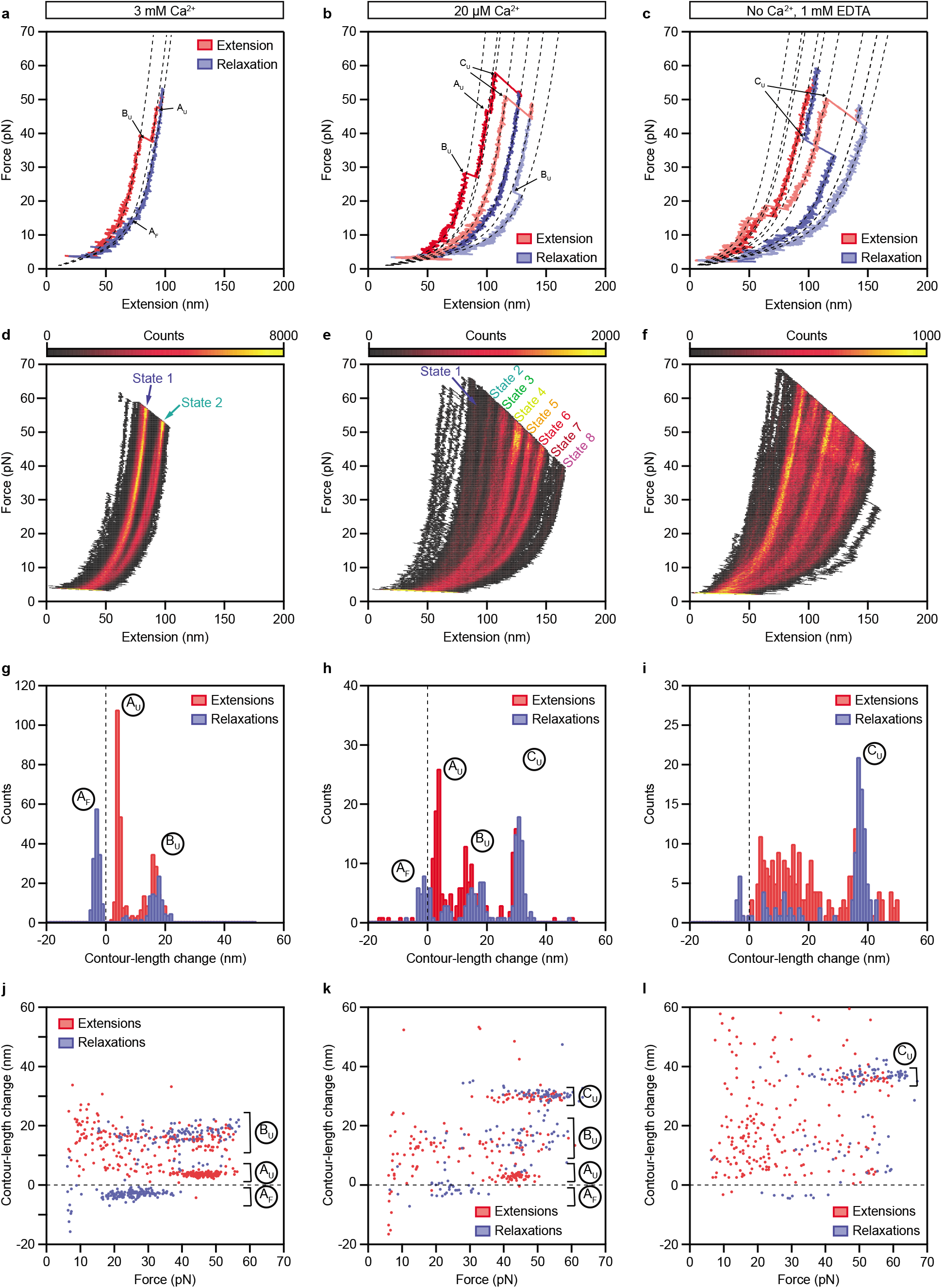
Force-extension measurements of PCDH15 monomers. **a**, At a Ca^2+^ concentration of 3 mM, individual force-extension cycles show two distinct classes of abrupt elongations, the unfolding events A_U_ and B_U_, as well as refolding events of class A_F_. The dashed lines represent fits to the trajectories by a protein model. **b**, Reducing the Ca^2+^ concentration to 20 μM elicits an additional class of unfolding events, C_U_, corresponding to the unfolding of entire cadherin domains. **c**, In the absence of Ca^2+^, unclassifiable structural changes occur in conjunction with the well-defined events C_U_. **d-f**, A heatmap displays all the force-extension cycles for a single representative molecule at each Ca^2+^ concentration. The data were binned into pixels of 1 nm x 0.1 pN. A much smaller portion of the state space is accessible for 3 mM [Ca^2+^] than for 20 μM [Ca^2+^] or in the absence of Ca^2+^. **g**-**i**, Histograms of the contour-length changes of all abrupt elongations verify that these rips can be grouped into classes A_F_, A_U_, B_U_, and C_U_ at Ca^2+^ concentrations of 3 mM and 20 μM. In the absence of Ca^2+^, most of the contour-length changes are more broadly distributed. **j-l**, Plots of the contour-length change of every rip against the force at which that event occurred reveal the force distributions of each class of structural change. Note that the extensions never completely unfolded a PCDH15 molecule, so elongations could occur even during the relaxation phases. All force-extension cycles were sampled at intervals of 10 μs and smoothed to a temporal resolution of 1 ms. The waiting times between cycles were 0.2 s for 3 mM [Ca^2+^], 2 s for 20 μM [Ca^2+^], and 4 s in the absence of Ca^2+^. The number of cycles recorded was 500 for 3 mM [Ca^2+^] and 200 for 20 μM [Ca^2+^] and in the absence of Ca^2+^.

## Conformational changes

Rips in the force-extension relations at physiological forces suggest that PCDH15 exists in different structural states during hearing. At a saturating Ca^2+^ concentration, the conformational states accessible to the protein are limited: the state-space heatmap reveals only two major configurations (Figure 2d). The second of these is further divided into two sub-states separated by a difference of only a few nanometers in contour length. State 1 reflects the extensibility of fully folded PCDH15 (Extended Data Note 1). By fitting a polymer model to the force-extension relations for five molecules, we find that State 2 arises from a combination of two classes of conformational changes leading to mean elongations of 4.0 ± 0.2 nm and 15.8 ± 0.7 nm (means ± SEMs; respectively distributions of A_U_ and B_U_ in Figure 2g; Table 1). The structural origin of these conformational changes is unknown and difficult to determine owing to the large size of the protein. We can, however, rule out the unfolding of entire cadherin domains as the origin of the unfolding events. The length of the folded peptide in each of the eleven cadherin domains ranges from 94 to 123 amino acids, with a mean of 104 residues. At a contour length of 0.39 nm per residue^31^, the unfolding of each cadherin domain is expected to augment the contour length by about 36 nm: an elongation of 40.6 nm less 4.5 nm to account for the loss of the folded cadherin domain. The observed contour-length changes of the elongations A_U_ and B_U_ therefore represent protein rearrangements less extreme than the unfolding of a cadherin domain. At a saturating Ca^2+^ concentration and physiological forces, we never observed length changes in the wild-type protein great enough to account for the unfolding of entire cadherin domains.

**Table 1.** Material property values of PCDH15.

| [Ca<sup>2+</sup>] | *b* (nm) | *l*<sub>Clinker</sub> (nm) | *k*<sub>folded</sub> (mN/m) | Δ*l*<sub>C<sub>A</sub></sub> (nm) | Δ*l*<sub>C<sub>B</sub></sub> (nm) | Δ*l*<sub>C<sub>C</sub></sub> (nm) |
| --- | --- | --- | --- | --- | --- | --- |
| 3 mM | 2.9 ± 0.5 | 1.4 ± 0.5 | 9 ± 4 | 4.0 ± 0.2 | 15.8 ± 0.7 | — |
| 20 μM | 3.1 ± 0.4 | 1.3 ± 0.5 | 11 ± 5 | 3.9 ± 0.4 | 15.0 ± 0.6 | 35 ± 1 |

Measurements are given as means ± SEMs for five experiments with 3 mM [Ca<sup>2+</sup>] and for eight experiments with 20 μM [Ca<sup>2+</sup>].

We next asked whether there are unique structural features in PCDH15 that give rise to the elongations A_U_ and B_U_, or whether several different conformational changes, each with a similar contour-length change, underlay the observed distributions. In most of the extension-relaxation cycles we did not observe more than one of either class of events (Extended Data Fig. 2). The rare occasions in which several events A_U_ or B_U_ were detected in a single trace were not reproducible across proteins or trials and could thus be explained as the rupture of non-specific interactions between the protein and either of the confining surfaces. We conclude that a single, unique structural alteration of PCDH15 is responsible for event A_U_, whereas a distinct structural change results in event B_U_, precluding the occurrence of several events of either type within the same cycle. Interestingly, the force sensitivity of events A_U_ is much narrower than that of events B_U_ (Figure 2j). From these distributions we determined the statistical dependence of both classes: an event A_U_ generally follows an event B_U_ (*p* < 0.05 and *p* < 0.1 for respectively three and two of the five tested molecules). It is plausible that both structural changes resulted from the same cadherin domain, with elongation B_U_ leading to a destabilization that facilitated elongation A_U_.

We found the protein in State 1 at the beginning of many extension-relaxation cycles and concluded that there is a high probability of refolding of both event A_U_ and event B_U_ between cycles. We indeed routinely detected refolding events A_F_ during the relaxation phase of our protocol (Figure 2g), but rarely observed refolding events B_F_. The latter events probably occurred only at very low forces, for which the slight shortening was lost in Brownian noise (Extended Data Fig. 3).

We next reduced the Ca^2+^ concentration to the physiological value of 20 μM and exposed a tethered PCDH15 protein to the same force protocol. The extension-relaxation cycles showed conformational changes identical to the previously observed classes (Figure 2h; A_U_: 3.9 ± 0.4 nm, B_U_: 15.0 ± 0.6 nm, means ± SEMs, *N* = 8). At this Ca^2+^ concentration, however, an additional class of unfolding events emerged with a contour-length change of 35 ± 1 nm (mean ± SEM; C_U_ in Figure 2b,h), in excellent agreement with the elongation expected for unfolding of an entire cadherin domain^32^. At holding forces of 2-4 pN the refolding of cadherin domains was a slow process and occurred on a time scale of seconds, in line with other proteins that feature immunoglobulin-like motifs^33^. In some extension-relaxation cycles we observed the successive unfolding of several cadherin domains (Extended Data Fig. 4). Because unfolding of any of the eleven extracellular cadherin domains should increase the contour length by a similar amount, we could neither assign unfolding events to particular domains nor elucidate the sequence in which the domains unraveled.

The heatmap of all force-extension relations originated from a mixture of the unfolding events A_U_, B_U_, and C_U_ and their respective refolding events. The contour lengths that gave rise to the annotated States 1 to 8 (Figure 2e) were a consequence of the unfolding of up to three cadherin domains in series with up to one unfolding event of type B_U_. Structural changes of type A_U_ and A_F_, which are clearly visible in the heatmap for a Ca^2+^ concentration of 3 mM as a subdivision of State 2 (Figure 2d), are not apparent in the state space at 20 μM [Ca^2+^]. It is possible that there were Ca^2+^-dependent sub-nanometer changes in the contour length that averaged out small effects of events A_U_ and A_F_ in the heatmap. We again investigated a potential sequential dependence of unfolding classes by their force distributions (Figure 2k): even though events of type A_U_ followed events of type B_U_ at 3 mM [Ca^2+^], at a physiological Ca^2+^ concentration these structural changes in eight molecules were independent of one another (*p* > 0.1 for each molecule). Moreover, events of types A_U_ and B_U_ were also independent of events of type C_U_, the unfolding of entire cadherin domains.

We next investigated the mechanics of PCDH15 in the absence of Ca^2+^. Representative force-extension relations feature a plethora of conformational changes (Figure 2c,i), many of which could no longer be clearly grouped into any of the classes A_U,F_ and B_U_. Events with a mean of 37 ± 2 nm (mean ± SEM for five molecules) continued to characterize a well-defined class C_U_. The heatmap of all extension-relaxation cycles had a structure reminiscent of that at a Ca^2+^ concentration of 20 μM (Figure 2f). The structure in the absence of Ca^2+^ arose from the unfolding of a discrete number of cadherin domains in series with the unclassifiable shorter structural changes that likely represented the partial unfolding of one or more domains. This lack of well-defined short structural changes was also evident from the force distribution of the observed rips (Figure 2l).

## Stiffness of PCDH15

It is unknown whether tip-link cadherins are completely or only partially folded during normal hearing. We therefore investigated the stiffness not only of folded PCDH15 but also that of conformational states with a progressively greater number of unfolded domains. The total stiffness of PCDH15 comprises both enthalpic and entropic components, whose contributions we quantified by fitting the force-extension relations with a model of the protein as a freely jointed chain^34^ formed by the eleven folded cadherin domains in series with a worm-like chain^35^ representing the ten unstructured linker regions. We allowed enthalpic extensibility through a Hookean spring constant and included an additional worm-like chain to model any unfolded portions of the protein (Extended Data Fig. 5). Because the unfolded polypeptide chains and the linker regions are structurally similar, we modeled them both with the same persistence length; fits to the data for thirteen molecules yielded *lp*_peptide_ = 0.49 ± 0.04 nm (mean ± SEM). For folded PCDH15 at a Ca^2+^ concentration of 3 mM, we found a length of 2.9 ± 0.5 nm for each of the eleven solid segments of the chain, a length of 1.4 ± 0.5 nm for each of the ten flexible linkers between the solid segments, and an enthalpic spring constant of 9 ± 4 mN·m^-1^ for the Hookean stiffness of the protein (means ± SEMs, *N* = 5 molecules, Table 1). The full length of a solid segment combined with its associated linker region was 4.3 ± 0.7 nm, in excellent agreement with the value of 4.5 nm per cadherin repeat from crystal structures of cadherin domains^23^. Much to our surprise, these values did not change when the Ca^2+^ concentration was lowered to 20 μM, the physiological level in the cochlea: an elevated Ca^2+^ concentration stabilizes cadherin domains against unfolding but does not augment the stiffness of the folded protein. The stiffness predicted by our model is in good agreement with the slopes of the different states in the state-space heatmap (Figure 3a,b).

**Fig. 3.**
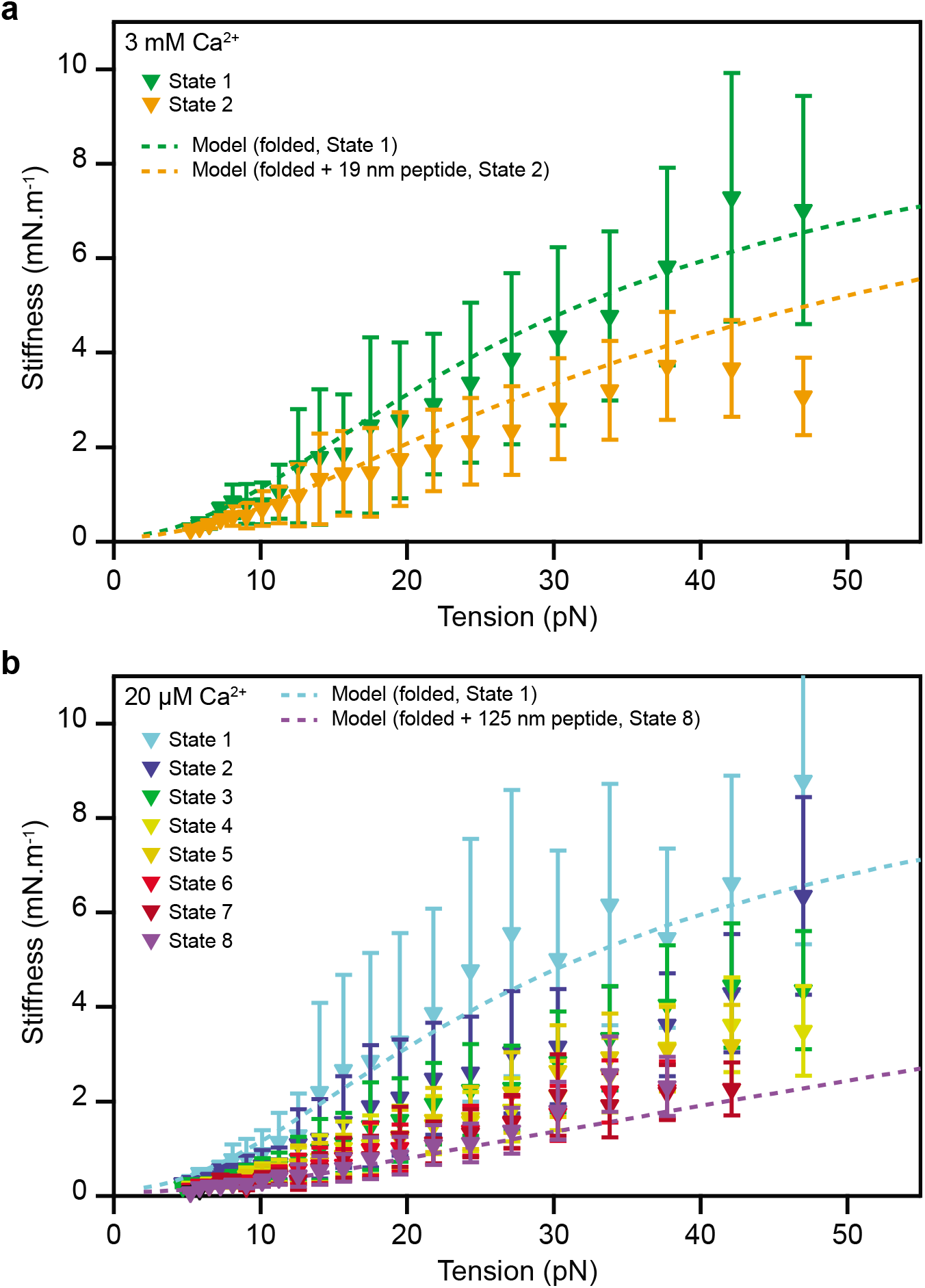
Stiffness of monomeric PCDH15. **a**, The stiffnesses of the different conformational states of PCDH15 at a Ca^2+^ concentration of 3 mM correspond to the slopes of the highly occupied regions of the state space in Figure 2d,e and are corrected for the stiffness of the molecular tags and anchors. The green dashed line represents the stiffness of our model of State 1, the fully folded protein, with the parameter values of Table 1 (*b* = 3.0 nm, *l*c_linker_ = 1.35 nm, *k*_folded_ = 10 mN·m^-1^; parameter values were averaged over both Ca^2+^ concentrations). The orange dashed line represents the model for State 2, with an additional 19 nm segment of unfolded protein with a persistence length of 0.49 nm representing the combined effect of the events A_U_ and B_U_. **b**, The corresponding data for a Ca^2+^ concentration of 20 μM capture a variety of unfolding events leading to States 2-8. The light-blue dashed line represents the fully folded protein (State 1); the purple dashed line depicts the stiffness of the protein in State 8, with an unstructured peptide 125 nm in length to represent the unfolding of three cadherin domains in series with contour-length changes of 15 nm and 4 nm. The experimental data are means ± SEMs for five molecules and six molecules at a Ca^2+^ concentration of respectively 3 mM and 20 μM.

Across all states and Ca^2+^ concentrations the measured and predicted stiffness of the protein is much smaller than its enthalpic stiffness of about 10 mN·m^-1^. The additional compliance is entropic, arising from the thermal motion of the individual cadherin domains and from thermal undulations in the inter-domain linker regions and unfolded polypeptide chains. When PCDH15 is tensed, this thermal kinking is smoothed out and the protein elongates. The progressive unfolding of domains further softens the protein by introducing additional disordered polypeptide chains (Figure 3b). At high forces most thermal bends have been straightened and the enthalpic elasticity begins to dominate the protein’s response. Importantly, we find that for physiological tension the protein’s response to force is dominated by entropic elasticity. The protein’s stiffness approaches its enthalpic value only for unphysiologically high tensions (Figure 3, Extended Data Fig. 6).

## Unfolding of cadherin domains under forces relevant for hearing

Elevated tension not only increases the stiffness of PCDH15 but also heightens the likelihood that entire cadherin domains unfold. Do cadherin domains unfold during normal hearing? If so, do they refold under physiological conditions, or could tip links with persistently unstructured regions exist *in vivo*?

To determine the unfolding rate of cadherin domains under physiological tensions, we transformed the force distributions of type C_U_ domain-unfolding events into unfolding rates as a function of constant force (Figure 4)^28^. For a given unfolding event we could not determine which of the eleven cadherin domains had unfolded, so our result was an average over several or all of the domains. The transformation additionally assumed that there was no cooperativity between the unfolding of individual domains. If the unfolding of one domain were to increase the probability that an adjacent domain would unravel, for example, our computed unfolding rates would have systematically overestimated the rate at which fully folded tip links unfold.

**Fig. 4.**
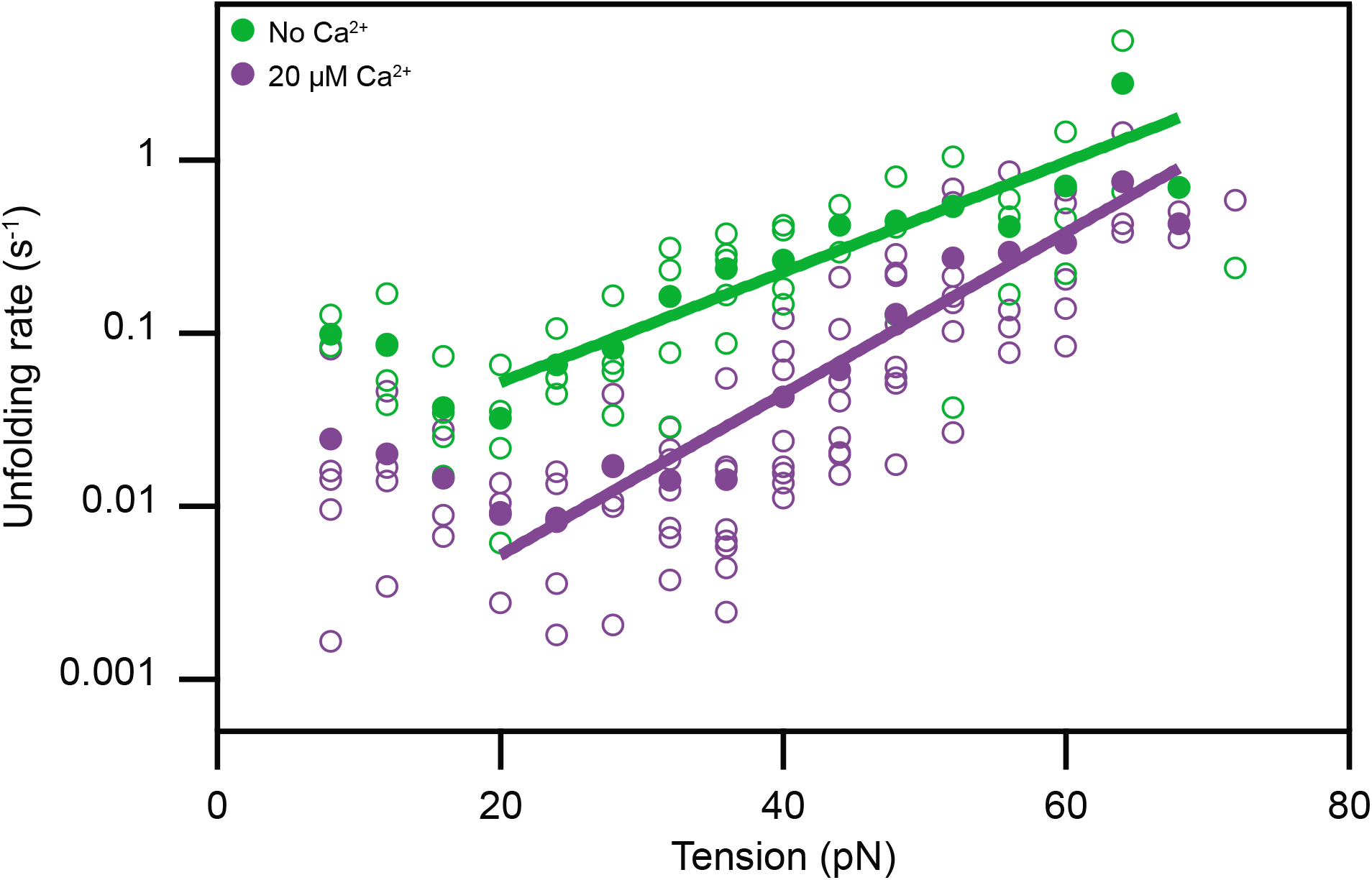
Tension-dependent unfolding rates of a single cadherin domain. **a**, Assuming that all domains of PCDH15 are similar and unfold independently, we estimate the rate at which individual EC domains unfold as a function of tension. Domains unfold much more readily in the absence of Ca^2+^ (green) than at a physiological Ca^2+^ concentration of 20 μM (purple). The filled circles represent the means for eight molecules at 20 μM [Ca^2+^] and for five molecules in the absence of Ca^2+^. The outlined circles, which represent the data for individual molecules, provide an estimate of the data’s spread. The solid lines are fits of Bells’ model^44^ to the data: for 20 μM Ca^2+^, the unfolding rate at zero force is *k*_0_ = 0.0006 ± 0.0002 s^-1^ and the transition-state distance is *x*^‡^ = 0.44 ± 0.04 nm; in the absence of Ca^2+^, *k*_0_ = 0.012 ± 0.006 s^-1^ and *x*^‡^ = 0.30 ± 0.04 nm.

We computed unfolding rates for Ca^2+^ concentrations of both 20 μM and zero (Figure 4). As expected, the presence of Ca^2+^ stabilized cadherin against force-induced unfolding: in the ion’s absence the domains unfolded many times faster than in its presence. With a physiological Ca^2+^ concentration and at 20 pN of tension, the upper range of physiological resting values, a single cadherin domain unfolded at a rate of approximately once every 100 s. A fully folded PCDH molecule—consisting of 11 domains—then unfolded a domain every 10 s. At even higher tensions this rate rapidly increased, to roughly 0.4 s^-1^ for the unfolding of a single cadherin domain at a tension of 60 pN. Because unfolding events at forces of 10-20 pN were extremely rare and might have corresponded to transitions from molten-globule states (Extended Data Note 2), we were unable to reliably calculate force-dependent unfolding rates for this force range. Extrapolation of the available data nevertheless suggests rates of approximately 0.003 s^-1^ at 10 pN of tension. These results indicate that unfolding of cadherin domains would not take place within individual cycles of an auditory stimulus: for physiological forces the unfolding rate is much too low to follow stimuli with frequencies ranging from hundreds to thousands of hertz. However, provided that at the link’s resting tension a domain’s unfolding rate exceeds its refolding rate, cadherin domains in a tensed tip link could exist in a permanently unfolded state. We never observed refolding of cadherin domains during any of the recorded extension-relaxation cycles, even though such events should be easily detectable at tensions exceeding 4 pN (Extended Data Fig. 3). Refolding instead occurred only during the holding phase between successive cycles, provided the holding force was below 4 pN and the waiting time was on the order of several seconds. We conclude that for tensions higher than 20 pN per molecule the unfolding rate—although very small—exceeds the refolding rate. Our data indicate that this is also the case for the force range of 4-20 pN, but owing to the possible influence of molten-globule states we could not determine this with certainty. Our result suggests that some cadherin domains in tip links *in vivo* exist in a perpetually unfolded state. Such unfolded states would decrease the protein’s stiffness (Figure 3b) and could be a mechanism by which a tip link softens even under high tension.

The critical force at which the unfolding and refolding rates of a cadherin domain are equal remains to be determined. However, the giant muscle protein titin, which has immunoglobin folds similar to those of cadherin, exhibits a critical force^33^ of 5.4 pN. If cadherin domains feature a similar value, the resting tensions in low-frequency hair cells might be less than the critical force and bias the tip link’s domains towards a fully folded state, whereas the tip link might occur in a partially unfolded state in hair cells sensitive to high frequencies.

## Effect of a mutation associated with hearing loss

Over one hundred mutations of the tip-link cadherins cause hearing loss in humans^36^. The deletion of residue V767 in EC7 is particularly interesting, for it leads to deafness—stemming from a deficit in the cochlea, with its low Ca^2+^ concentration—but not a loss of function in the vestibular labyrinth—which enjoys a higher Ca^2+^ concentration^37^. This mutation evidently does not hinder tip-link formation, but might change the elastic properties of the link.

We investigated how this deletion affects the mechanical properties of monomeric PCDH15 (V250D, ΔV767) and found that there was a small but detectable probability that force unfolded a complete cadherin domain even when PCDH15 was saturated with Ca^2+^ (Figure 5). During identical treatment of Ca^2+^-saturated proteins without pathologic mutations we never observed the unfolding of complete domains (Figure 2g). By shortening one strand of EC7, the mutation likely caused a slight misalignment of amino-acid residues and thus destabilized the domain. When we performed experiments at the physiological Ca^2+^ concentration of 20 μM, we could not detect a difference in domain unfolding between the mutant and wild-type proteins.

**Fig. 5.**
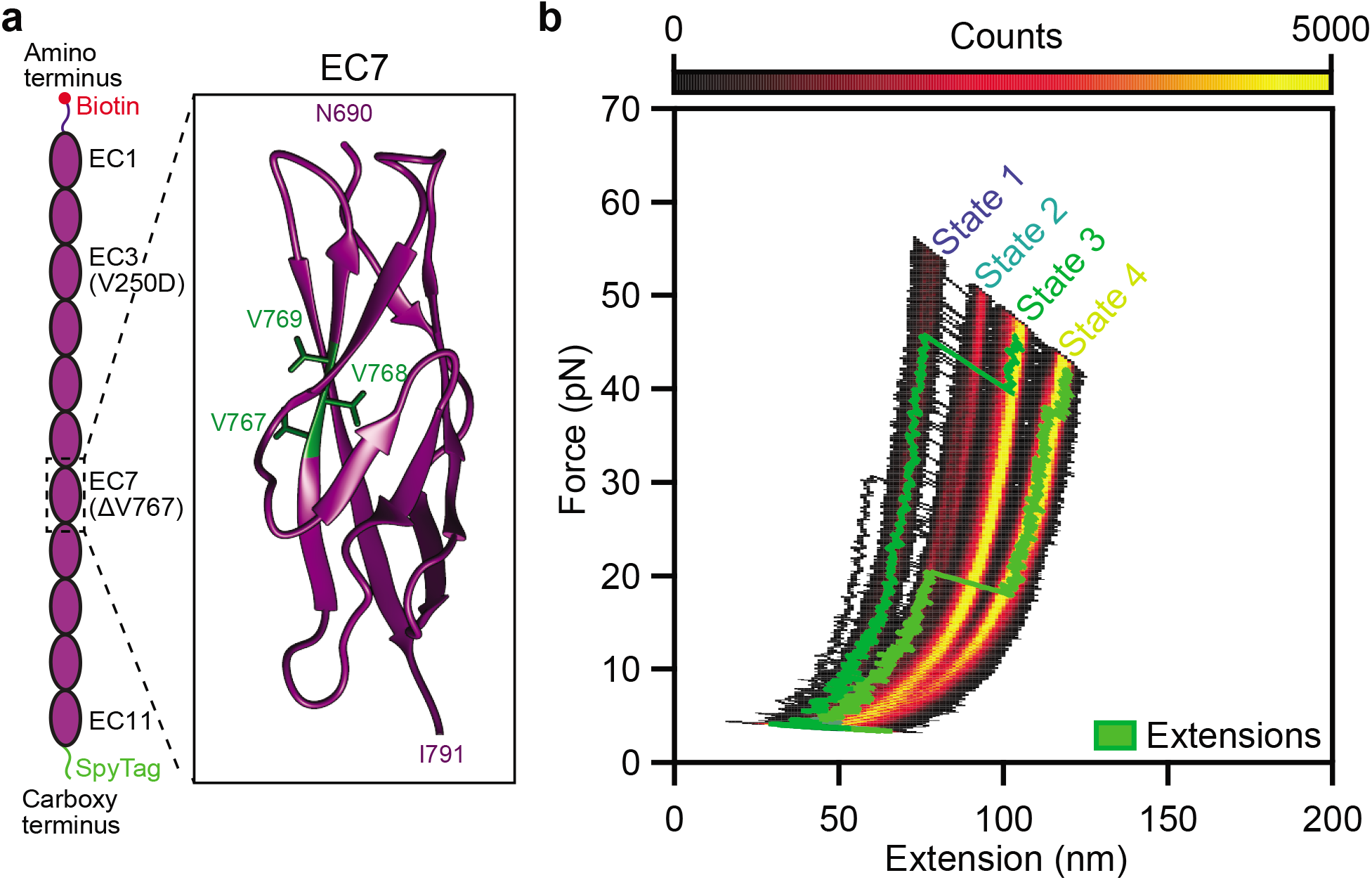
Effect of a hearing loss-associated mutation on PCDH15 mechanics. **a**, We deleted valine 767 in the seventh EC domain of PCDH15. As indicated in the crystal structure (PDB ID code 5W1D, image generated with UCSF Chimera), V767 is located in the F strand of the cadherin fold. **b**, A state-space heatmap for 500 extension-relaxation cycles reveals that at 3 mM [Ca^2+^] the mutant protein can assume four distinct conformational states. The two additional states not observed in the wild-type protein result from unfolding of the pathologic cadherin domain in series with the usual States 1 and 2. Unfolding of the pathological domain is rare and occurs in only a few cycles, two of which are superimposed upon the heat map (green traces). The waiting time between cycles was 0.2 s.

## Discussion

The behavior of a gated ion channel is usually binary: the channel is open or closed. A mechanically activated channel can nevertheless signal fine nuances of a stimulus by rapidly fluttering between the two states, such that the average open probability provides a smoothly graded representation of the stimulus. A gating spring makes this possible: tensed by a stimulus and battered by thermal noise, the spring continuously adjusts the open probability of the associated channel over a significant range of inputs. This range is determined by the gating spring’s stiffness, and thus by such molecular details as the entropic elasticity and folding transitions demonstrated here.

The stiffness of gating springs in outer hair cells increases with heightened resting tension along the tonotopic axis^16^, from 1.9 mN·m^-1^ at 7 pN of tension to 5.5 mN·m^-1^ for 50 pN. Simulations of short segments of tip-link proteins indicated that they are orders of magnitudes too stiff to account for these values^7,8^. However, our single-molecule experiments on the extracellular domain of PCDH15 reveal that the protein has a stiffness comparable to that of gating springs *in vivo* and displays similar strain-hardening. Across the physiological force range most of PCDH15’s compliance is of entropic origin; the protein’s enthalpic stiffness of 9 mN·m^-1^ emerges only at very high tension. Our stiffness values (Figure 3) are systematically lower than those found for gating springs *in vivo*, which is not surprising because our measurements tested only a monomer of one constituent protein. The dimeric arrangement of PCDH15 roughly doubles the enthalpic stiffness of the monomer. Moreover, a tip link adopts a helical structure^17,19^ that likely reduces the magnitude of its thermal undulations, decreasing the entropic contribution to the tip link’s mechanics and further increasing its stiffness. Finally, the arrangement of a dimer of CDH23 in series with the PCDH15 dimer is expected to reduce the stiffness by about 70 %. At very high tensions, when entropic effects are largely suppressed and enthalpy dominates, we estimate that the stiffness of the full-length, dimeric tip link is 6 mN·m^-1^ (Extended Data Note 3), a value in good agreement with the stiffness of strongly tensioned gating springs *in vivo*. These results suggest that the tip-link cadherins are a major component of the gating spring for mechanotransduction in hair cells.

The stiffness of hair bundles increases at low Ca^2+^ concentrations^38^, an observation that can now be interpreted as strain-hardening of the tip links. Low Ca^2+^ levels cause molecular motors to upregulate tip-link tension, which suppresses each link’s thermal motion and increases its stiffness. In addition to this indirect modulation of tip-link stiffness by the Ca^2+^ concentration, we found that Ca^2+^ also directly affects the rate at which cadherin domains unfold under force. At a tension of 20 pN and at a Ca^2+^ concentration of 20 μM, individual cadherin domains unfold an order of magnitude more slowly than in the absence of Ca^2+^. We never observed the unfolding of entire cadherin domains at a Ca^2+^ concentration of 3 mM. The sensitivity of PCDH15 to Ca^2+^ suggests that the variable concentrations of the ion in different receptor organs tunes the mechanical properties of tip links and thus of hair bundles to the organs’ specific requirements. Even within a single organ, the guinea pig’s cochlea, the Ca^2+^ concentration increases fourfold along the tonotopic axis from base to apex^26^. This gradient might adjust tip-link stiffness to accord with the frequency response of the individual hair cells. Finally, hair cells can enhance the local Ca^2+^ concentration around their hair bundles through the activity of membrane Ca^2+^ pumps^39^. This phenomenon raises the interesting possibility that the stiffness of tip links is modulated by the locally varying Ca^2+^ concentration in response to hair-cell activity.

We found that the elastic properties of folded PCDH15 are surprisingly independent of the Ca^2+^ concentration. This result seems to contradict the impression conveyed by electron-microscopic images^3^, in which cadherins transition from a disordered globular conformation to a rod-like chain of domains in the presence of progressively larger amounts of Ca^2+^. Disordered states should make larger entropic contributions to PCDH15’s elasticity than ordered states, a difference not apparent in our data. Note, however, that the divergence between electron-microscopic images and single-molecule data has also been observed for other proteins such as titin^40^. A possible explanation is that the configurations of proteins adsorbed to electron-microscopic substrates are far from their equilibrium conformations, so that the variations in shape do not accurately capture the thermal motion in solution^41^.

The force-extension relations of hair bundles reveal that, for unphysiologically large stimuli, gating springs can stretch^42^ by at least 120 nm, a value thought to be incompatible with the extensibility of tip-link cadherins. It has consequently been suggested that the gating spring’s stiffness stems from the elasticity of the plasma membrane or cytoskeleton into which the tip links insert^43^. We have shown that invoking such sources of elasticity is unnecessary: cadherin domains in the tip-link proteins can unfold under physiological stimuli, albeit at a low rate, and such unfolding events become very likely at high forces. Extension of the tip link by 120 nm is easily possible through the unfolding of several cadherin domains. In further support of domain unfolding, the length distribution of tip links in the bullfrog’s hair bundles, as determined by electron-microscopic tomography, features two distinct classes^13^ with means near 110 nm and 170 nm. The unfolding of two cadherin domains per tip-link monomer could account for this length difference. Such unfolding events could soften the gating spring at high resting tensions and protect both the tip link and the associated mechanotransduction machinery from damage during loud sounds.

In addition to the unfolding of entire cadherin domains, we also observed partial domain unfolding with contour length increases of 4 nm and 15 nm. Future single-molecule work will be necessary to elucidate the structural correlates of these conformational changes and to determine what role they play in hearing. Additional experiments will also be necessary to test the stiffness of PCDH15 dimers and of the full tip link and to confirm that domain unfolding occurs for those constructs and for tip links *in vivo*.

## Acknowledgments

The authors thank Julio Fernandez and Jaime Andrés Rivas-Pardo for guidance concerning the chemical functionalization of glass substrates; Brian Fabella and Vadim Sherman for high-precision fabrication; Brandon Razooky and Maria Vologodskaia for assistance with cloning; and Anna Kaczynska for help with protein expression. This work was partially supported by a Junior Fellow award from the Simons Foundation and a Pilot Grant from the Kavli Foundation to T.F.B. A.J.H. is an Investigator of Howard Hughes Medical Institute.

## Author contributions

T.F.B. and A.J.H designed the project. T.F.B., F.E.H., A.O., S.S.M., M.E.S., and U.M. created cDNA clones for PCDH15. G.D. and L.S. expressed and purified the recombinant protein. I.V.C., T.F.B., F.E.H., and A.O. prepared experimental substrates and samples. I.V.C. calibrated the Ca^2+^ concentrations. T.F.B. designed, built, developed software for, and characterized the photonic force microscope. T.F.B., F.E.H., and A.O. conducted the biophysical experiments. T.F.B. carried out the data fitting and analysis of statistical significance. T.F.B., I.V.C., F.E.H., A.O, and A.J.H. wrote the paper.

## Methods

### Molecular cloning

Plasmids were assembled by Gibson assembly in a one-step isothermal reaction using home-made master mixes^45^. We assembled in a pLEXm backbone^46^ a construct encoding the protein **signal peptide-QYDDDWQYED-Avitag-GSGSGS-PCDH15(EC1-11, V250D)-GSGSGS-Spytag-6xHis**. The deletion mutant (V250D, ΔV767) was assembled in a similar reaction. The PCDH15 sequence was isoform 1 from *Mus musculus* (UniProtKB entry Q99PJ1). The signal peptide comprises the native sequence that leads to secretion of PCDH15. Two tags for the site-specific confinement of PCDH15 were fused to the termini of the protein. The Avitag, with the sequence **GLNDIFEAQKIEWHE**, is recognized by a biotin ligase (BirA-500, Avidity, Aurora, CO, USA), which covalently biotinylates the lysine side chain. The SpyTag had the sequence **AHIVMVDAYKPTK**. The tags were attached to PCDH15 through flexible **GSGSGS** linkers.

### Expression of recombinant PCDH15

All constructs were transfected with 40 kDa polyethyleneimine (Polysciences, Inc., Warrington, PA, USA) into suspension-adapted HEK293 cells (Freestyle, R79007, Thermo Fisher Scientific, Waltham, MA, USA). Seven to nine days post transfection, the medium was collected and secreted proteins were purified by Ni^2+^-affinity chromatography. Proteins were further purified by size-exclusion chromatography (Superose 6 10/300 GL,17517201, GE Healthcare Bio-Sciences, Pittsburgh, PA, USA) in 10 mM Tris, 150 mM NaCl, and 3 mM CaCl2. The purified proteins were concentrated to 1.5 mg/ml and biotinylated for 1 hr at 30 °C with biotin ligase (BirA 500, Avidity, Aurora, CO, USA). The samples were then used immediately or mixed with equal volumes of glycerol and stored at −20 °C for up to four months.

### Design of the control linker peptide

In order to test the mechanical properties of the linkers and anchors of the assay in the absence of PCDH15, we fused the SpyTag and AviTag by a flexible **GGGSGGGS** linker to produce a control linker peptide with the sequence **AHIVMVDAYKPTKGGGSGGGSGLNDIFEAQKIEWHE** (Genscript, Piscataway, NJ, USA). After biotinylation, this peptide is capable of tethering streptavidin-coated probe beads to SpyCatcher molecules on pedestal beads, representing a control single-molecule assay that contains all components except for the PCDH15 protein.

### Site-specific attachment of PCDH15 through short molecular anchors

The carboxy terminus of PCDH15 was modified with a SpyTag capable of forming a covalent bond with a small, globular protein called SpyCatcher, which was linked to the surface of the pedestal bead. The protein’s amino terminus was biotinylated to allow its strong, site-specific attachment to streptavidin molecules on the probe bead (Figure 1d,f). The SpyCatcher-SpyTag complex is mechanically stable up to nanonewtowns of force, well beyond the range relevant for hearing^47^. The streptavidin-biotin interaction has a lifetime of about 30 s for the highest forces applied in this work^16,48^. These properties made this set of tags and binding partners ideal for the site-specific confinement of proteins under force.

### Conjugation of pedestal beads with SpyCatcher molecules

Cys-SpyCatcher molecules (EOX004, Kerafast, Boston, MA, USA) were conjugated to aminated silicon dioxide microspheres (140414-10, Corpuscular, Cold Spring, New York, USA) through short, bifunctional polyethylene glycol spacers. To deprotonate the surface amine groups, 200 μl of the beads was washed once and resuspended for 1 hr at room temperature in 100 μl of 50 mM sodium tetraborate buffer (11625, Sigma Aldrich, St Louis, MO, USA) at pH 8.5. Deprotonation is necessary for efficient covalent coupling of the *N*-hydroxysuccinimide-PEG_12_-maleimide spacer (22112, ThermoFisher, Waltham, Ma, USA), which we added to the beads to a final concentration of 50 mM of the linker and a final volume of 160 μl. The resulting mixture was incubated for 30 min at room temperature, washed three times with 1 ml of Hepes-buffered saline solution (HeBS; 20 mM Hepes and 100 mM NaCl), and after the final wash resuspended into 250 μl of HeBS.

Meanwhile, 0.5 mg of Cys-SpyCatcher protein was dissolved in 50 μl of HeBS and reduced for 1 hr with tris(2-carboxyethyl)phosphine (Immobilized TCEP Disulfide Reducing Gel, 77712, ThermoFisher Scientific, Waltham, MA, USA) according to the manufacturer’s instructions. The reduced SpyCatcher protein was then mixed with 55 μl of 100 mg/ml sulfhydryl-blocked bovine serum albumin (BSA; 100-10SB, Lee Biosolutions, Inc., Maryland Heights, MO, USA) in HeBS. The bead solution was injected into the protein mixture and incubated at 4 °C overnight to allow the covalent attachment of the SpyCatchers’ unique cysteines to the maleimide residues on the pedestal bead. The beads were then washed three times with 1 ml of HeBS and resuspended to a volume of 100 μl in HeBS. Any unreacted maleimide was quenched by addition of 100 μl of 1 M L-cysteine (11033-016, Gibco BRL, Gaithersburg, MD, USA) in HeBS and incubation for 1 hr. Finally, the beads were washed three times and stored at 4° C in HeBS with 0.02 % sodium azide (71289, Sigma-Aldrich, St. Louis, MO, USA).

### Covalent attachment of pedestal beads

We covalently attached pedestal beads through the surface amine groups of their SpyCatcher molecules to COOH-modified glass coverslips through carbodiimide crosslinking. Glass coverslips (12-545-81, Thermo Fisher Scientific, Waltham, MA, USA) were cleaned by sonication in ethanol for 15 min. After drying under a stream of oxygen, the coverslips were transferred to a solution of 1.5 g Nochromix (Godax Laboratories, Cabin John, MD, USA) in 60 ml sulfuric acid (A300S-500, Thermo Fisher Scientific, Waltham, MA, USA) and incubated for 3 hr. We then washed the coverslips three times with deionized water followed by three washes with ethanol. During each wash the coverslips were sonicated for five minutes in a bath sonicator. The slides were dried under a stream of oxygen gas and oxidized for 30 min in an ultraviolet ozone cleaner (PC440, Bioforce Nanosciences, Salt Lake City, UT, USA). In this step oxidation occurred on only one side of the coverslips, the “functional” side, which was used in all the subsequent steps.

We vapor-deposited an aminosilane layer onto the oxidized glass surfaces by placing the coverslips into a gas-tight glass container together with—but not submerged in—100 μl of (3-aminopropyl)trimethoxysilane (281778, Sigma Aldrich, Inc, St Louis, MO, USA) dissolved in 10 ml of toluene (T324-1, Thermo Fisher Scientific, Waltham, MA, USA)^49^. The container was incubated overnight at 80 °C. Following the vapor deposition, the coverslips were washed three times in ethanol, dried, and placed into a solution of 50 mg succinic anhydride (S7626, Sigma Aldrich, St Louis, MO, USA) dissolved in 1 ml dimethyl sulfoxide (D4540, Sigma Aldrich, St Louis, MO, USA) and left to incubate at room temperature for at least 3 hr. This step converted the vapor-deposited amine groups into carboxyl groups and rendered the surface suitable for coupling. The carboxylated slides were then rinsed three times with ethanol and either used immediately or stored in ethanol for up to 72 hr.

### Assembly of samples

To assemble a sample, a carboxylated coverslip was dried and secured with vacuum grease onto a metal washer, so that its functional side contacted the washer (Extended Data Fig. 7). (1-ethyl-3-(3-dimethylaminopropyl)carbodiimide and *N*-hydroxysulfosuccinimide (respectively 77149 and 24510, Thermo Fisher Scientific, Waltham, MA, USA) were equilibrated to room temperature before 10 mg of each reagent was dissolved in 1 ml of activation-buffer solution containing 10 mM NaCl and 1 mM 2-(*N*-morpholino)ethanesulfonic acid (M3671, Sigma Aldrich, St Louis, MO, USA) at pH 6. Of the resulting solution, 50 μl was pipetted onto the functional surface of the mounted slide and left to incubate at room temperature for exactly 30 min to activate the slides with amine-reactive hydroxysulfosuccinimide esters. To remove any excess reagents, we carefully washed the activated slide by pipetting 2 ml of Hepes-buffered saline solution with Ca^2+^ (HeBS-Ca, 20 mM Hepes, 100 mM NaCl, 1 mM CaCl_2_) onto the mounted coverslip. Immediately after removing the solution except for a thin film to keep the active surface from drying, we pipetted pedestal beads in HeBS-Ca onto the activated surface and allowed to react for 2 hr. Mounting a second coverslip on the top of the washer closed the sample chamber. Access ports in the washer allowed the exchange of solutions within the chamber. To reduce non-specific interactions of PCDH15 with the beads, we exchanged the fluid to blocking-buffer solution containing 10 mg/ml sulfhydryl-blocked bovine serum albumin (100-10SB, Lee Biosolutions, Inc., Maryland Heights, MO, USA), 150 mM NaCl, 20 mM Tris-HCl, and 3 mM CaCl_2_ at pH 8. The sample was stored overnight at 4 °C before addition of the PCDH15 molecules.

### Sample preparation

PCDH15 monomers were diluted into blocking-buffer solution and flushed into a sample chamber. The molecules were sufficiently dilute to ensure that any tether was likely with an average probability exceeding 90 % to represent a single molecule of PCDH15, rather than two or more at once (Extended Data Table 1). We incubated the sample for 1 hr at room temperature to allow the carboxy termini of the PCDH15 molecules, each fused to a SpyTag, to covalently bind to the SpyCatcher proteins on the surface of the pedestal beads. The biotinylated amino termini were then directed radially outward from the pedestal beads and thus available for coupling to streptavidin-coated probe beads. The sample was subsequently washed with copious amounts of blocking-buffer solution to remove any unbound PCDH15 monomers.

The blocking buffer was replaced with sample-buffer solution consisting of 20 mM Tris-HCl pH 7.5, 150 mM NaCl, and 10 mg/ml sulfhydryl-blocked bovine serum albumin (100-10SB, Lee Biosolutions, Inc., Maryland Heights, MO, USA). The solution contained probe beads (CP01004, Bangs Laboratories, Fishers, IN, USA); depending on the experiment, it included 3 mM CaCl_2_, 20 μM CaCl_2_, or 1 mM EDTA. To protect PCDH15 from photodamage, we employed an oxygen-scavenging system consisting of 18 mM D-glucose (G-5400, Sigma-Aldrich, St. Louis, MO, USA), 13 U/ml pyranose oxidase (P4234, Sigma Aldrich, St Louis, MO, USA), and 8500 U/ml catalase (219261, Millipore Sigma, Burlington, MA, USA)^50,51^.

### Calibration of Ca2+ concentration

We used a fluorescence assay to confirm that the binding of Ca^2+^ to BSA does not significantly alter the concentration of free Ca^2+^ in sample-buffer solution (Extended Data Fig. 8). Using 3 μM of the fluorescent calcium indicator Fluo-5N (F14203, ThermoFisher Scientific, Waltham, MA, USA), we tested the fluorescence of the solution with and without 10 mg/ml BSA at various total Ca^2+^ concentrations. Our data show that BSA at this concentration does not noticeably change the concentration of free Ca^2+^, whereas a very high BSA concentration, 100 mg/ml, sequesters a significant amount of Ca^2+^ (Extended Data Fig. 8b).

### High-bandwidth and high-precision optical trapping and tracking

All data were acquired using a custom-built photonic-force microscope^22^, which in this instance was upright rather than inverted. The microscope could track the three-dimensional position of a weakly optically trapped, 1 μm-diameter probe bead with an integration time of 1 μs, sampled at a frequency of 100 kHz, with sub-nanometer precision. In brief, the position-sensing 1064 nm laser beam (Mephisto 500 mW, Coherent, CA, USA) was expanded and focused into the sample chamber through a high-numerical aperture water-immersion objective lens (UPlanSApo 60xW, Olympus, Tokyo, Japan). This beam formed a weak optical trap that confined the probe bead. We collected light forward-scattered by the probe bead together with the unscattered portion of the beam on a quadrant photodiode, where the two waves interfered. The signals of the four quadrants were related to the three-dimensional position of the bead in the optical trap^21^. The microscope’s position error over one extension-relaxation cycle with a duration of 1 s was given by its position noise from 1 Hz to 1 MHz, for which we measured root-mean-square values of 0.6 nm, 0.3 nm, and 0.4 nm along the x-, y-, and z-axes respectively^52,53^. Experiments were performed with the protein tether oriented along the y-axis, that with highest precision. Typical spring constants of the weak position-sensing trap were 6 μN·m^-1^, 7.5 μN·m^-1^, and 2 μN·m^-1^ along the x-, y-, and z-axes respectively. At very high stimulus forces of tens of piconewtons the probe bead was at its maximum extension of about 150 nm along the y-axis from the center of the position-sensing optical trap, corresponding to a maximum force generated by this trap of 1.1 pN. For lower stimulus forces and smaller extensions, the force was considerably less than 1 pN. Because the force generated by the position-sensing optical trap was very small compared to the stimulus force, we disregarded it in our analysis.

To record hundreds of cycles, we required observation times greater than a few seconds and therefore had to compensate the microscope’s slight mechanical drift. The position of the sample relative to the optical traps was controlled by a nano-positioning stage (Nano-View/M375HS, Mad City Labs, WI, USA), whose position we adjusted to compensate for the drift (see below). With drift compensation in effect, the root-mean-square deviation measured along the y-axis between DC and 1 MHz during 5 min of observation was 2 nm.

In addition to the previously described position-sensing weak optical trap, we added a second optical trap to the system to apply force stimuli to the tethered proteins. We chose a wavelength of 852 nm (DL852-500, Crystalaser, Reno, NV, USA), which is near a local minimum of the action spectrum of photodamage to biological material^54^. Using a beam-steering lens mounted on a three-dimensional piezoelectric-block translator (P.282.30, Physik Instrumente, Auburn, MA, USA), we could shift the position of the stimulus trap with respect to the position-sensing trap with nanometer precision. Over 18 min of observation time, the average drift of the two optical traps relative to each other was 3 pm·s^-1^. The 852 nm laser beam traversed an electro-optical modulator (LM13, Excelitas Technologies, Fremont, CA, USA) placed between crossed polarizing beam-splitting cubes, which allowed us to modulate the intensity and thus the stiffness of the stimulus trap.

### Drift compensation

Although we designed the microscope’s frame to minimize thermal drifts of the sample chamber with respect to the optical traps, a small drift of about 250 pm·s^-1^ remained^22^. To eliminate this drift, we implemented an active-feedback mechanism by using as a fiducial marker a second pedestal bead situated tens of micrometers from the pedestal to which a single molecule was tethered. The motion of the pedestal bead reflects the drift of the sample chamber with respect to the microscope frame. A camera (pco.edge 5.5, PCO, Kelheim, Germany) tracked the three-dimensional position of this pedestal bead at a frequency of 5 Hz. The pedestal’s position signal was then used as the input of a proportional-integral-differential feedback loop that adjusted the position of the nano-positioning stage to compensate for the sample’s drift. To test the fidelity of this method, we immobilized a probe bead on a sample chamber’s coverslip, positioned it at the focus of the position-sensing optical trap, and used the microscope to determine its position while drift compensation was active. We found that linear drifts were eliminated and that the root-mean-square variation of the position signal along the y-axis of 2 nm over 5 min remained.

### Calibration, linearization, and correction of the probe-position signal

The three-dimensional detector’s non-linear response was linearized and calibrated *in situ* for each individual probe bead^55^. The calibration depended upon the viscosity of the buffer solution, which we corrected for the presence of 10 mg/ml bovine serum albumin as described^56^.

The probe bead forward-scattered a portion of the position-sensing laser beam, and we detected the bead’s three-dimensional position by monitoring the interference pattern of the beam’s scattered and unscattered light wave on a quadrant photodiode^21^. Scattering of a small amount of the beam by the pedestal bead produced an artifact in the signal^22^ that was eliminated by the subtraction of a reference signal.

### Calibration and correction of the stimulus trap

Before each single-molecule experiment and for each individual probe bead, we measured the relation between the intensity of the stimulus laser and the stiffness of the stimulus trap. While monitoring the Brownian motion of the trapped probe bead with the position-sensing optical trap, we increased the power of the stimulus beam in 10 % increments. Fitting of the position signal’s power spectral density for each intensity with a hydrodynamically correct theory^57^ yielded the stiffness as a linear function of intensity. During our experiments, we sampled the intensity of the stimulus laser—and hence the trap’s stiffness—with the same sampling rate as the position signal of the probe bead.

We adjusted the intensity of the stimulus laser using an electro-optical modulator. At high attenuation, close to extinction, the beam profile at the exit of the modulator deviated from a Gaussian function, which laterally shifted the position of the stimulus trap within the sample chamber by a few nanometers. We recorded this shift during the calibration procedure for each probe bead and accounted for it when calculating the laser intensity-dependent force on the probe bead.

### Initiation of single-molecule experiments

To initiate a single-molecule experiment, we optically trapped a probe bead deep in solution and calibrated the position sensor and the stiffnesses of both optical traps. We next positioned the bead’s center at a height of 1 μm from the functionalized coverslip, with its mean axial position at the equator of the attached pedestal beads. This height was determined by slowly moving the coverslip towards the optical trap until the probe bead’s axial thermal motion began to be confined by the coverslip, then retracting the coverslip by an appropriate distance^52^.

The sample was moved laterally so that the probe bead was aligned along the y-axis with a pedestal bead. After recording a reference signal to account for light scattered by the pedestal we gently maneuvered the pedestal bead towards the optical trap until the probe’s Brownian motion along the y-axis began to be restricted by the pedestal. When a PCDH15 molecule was present on the surface of the pedestal bead and within reach of the Brownian motion of the probe bead, a single molecule tether formed between the amino terminal biotin on the protein and a streptavidin molecule on the probe bead. The concentration of PCDH15 was titrated so that only a fraction of such approaches resulted in tether formation, on average giving rise to a 90 % confidence of single-molecule conditions^58^ (Extended Data Table 1). The position of the stimulus trap was then displaced by 200 nm along the y-axis. Before the force was lowered to the holding level, a brief increase in the intensity of the stimulus beam provided a force pulse of several tens of piconewtons to the tethered protein. This operation ensured that no portion of the protein was nonspecifically attached to either of the confining surfaces and that the full contour of the protein linked the two beads at the beginning of the extension-relaxation cycles. Force ramps were then applied to the tethered protein by repeatedly increasing and decreasing the intensity of the stimulus trap (Extended Data Fig. 1).

### Control of non-specific attachments

For a successful single-molecule experiment it is imperative that the overwhelming majority of tethers between pedestal and probe beads constitutes PCDH15 molecules anchored at their amino and carboxy termini by respectively biotin and SpyTag: only a very small number of non-specific tethers should occur.

We tested whether streptavidin-coated probe beads would tether to SpyCatcher molecules on pedestal beads in the absence of PCDH15. Out of 65 attempts of initiating such non-specific tethering with two samples and four probe beads, only one bond formed, which ripped off immediately upon application of a stimulus force. We concluded that all stable tethers that we observed in our single-molecule experiments resulted from PCDH15 molecules or, in the case of control experiments to test the assay’s mechanics, from linker-peptide constructs.

To exclude any non-specific interactions between PCDH15 and either of the beads, we next confirmed that these PCDH15 tethers formed only if both the SpyTag-SpyCatcher and biotin-streptavidin interactions were present. Pedestal beads were coupled to a coverslip and incubated with 0.15 mg/ml PCDH15 in blocking-buffer solution for 1 hr at room temperature to allow the proteins to react with the pedestals. The coverslip was then washed with copious amounts of blocking-buffer solution to remove any free PCDH15 molecules before the addition of a high concentration of probe beads in blocking-buffer solution. The probe beads were allowed to bind to the PCDH15 molecules on the pedestal beads for 1 hr before the coverslip was washed once more and then imaged.

As a positive control, with both sets of anchors intact, we found an average of 3.25 probe beads bound to each pedestal bead for the given reaction conditions (Extended Data Table 2). To test whether the carboxy-terminal SpyCatcher-SpyTag anchor participated in the tether, we generated pedestal beads without SpyCatcher and attempted to attach probe beads to them through PCDH15 molecules using the procedure described above. We found that tethering of probe beads was completely abolished in the absence of SpyCatcher, confirming that the SpyCatcher-SpyTag interaction was an essential part of the formed tethers (Extended Data Table 2).

To determine whether the amino termini of our tethers were anchored to the probe beads through the biotin-streptavidin interaction, we attempted to tether probe beads to SpyCatcher-positive pedestals through PCDH15 molecules that had not been biotinylated. We again found that tether formation was completely abolished, confirming that the amino-terminal confinement in our single-molecule experiments occurred through the biotin-streptavidin interaction (Extended Data Table 2).

We concluded that our single-molecule assay was highly specific: molecular tethers formed in the presence of an appropriately tagged construct. If either of the pairs of anchors was disrupted, tether formation was completely abolished.

### Determination of the molecule’s anchor position

The anchor position of the protein tether along the axis of extension was an important parameter that had to be determined before our protein model could be fit to the data. We determined this anchor position by analyzing the three-dimensional probability distribution of the motion of the tethered probe bead in the absence of externally applied tension.The pedestal bead appeared as a forbidden volume in the spatial probability distribution of the tethered probe bead, a so-called three-dimensional thermal-noise image^22^. The intersection of the surface of the pedestal bead with the axis of extension was defined as the anchor position of the tether.

### Sources of uncertainty

Our data are subject to several different sources of measurement uncertainty. In the following we refer to the variability within one single-molecule experiments as “precision,” whereas we use “accuracy” to refer to the uncertainty between experiments.

The extension of a molecular tether could be measured with the same precision with which the photonic-force microscope could measure the position of the probe bead. To determine this value, we attached a probe bead to a glass coverslip, positioned it in the center of the position-sensing optical trap, and activated the microscope’s drift compensation. Between 1 Hz and 1 MHz, the interferometric position signal of the immobilized probe had a band-limited standard deviation of 0.3 nm along the y-axis, the axis along which we extended single-molecule tethers (Extended Data Table 3a). Over 5 min of observation, the standard deviation for the full bandwidth of DC to 1 MHz was 2 nm (Extended Data Table 3b).

The accuracy of the position sensor depends on the fidelity of its calibration, which can be tested by its comparison to a calibrated standard. We confined a probe bead in the weak position-sensing optical trap, calibrated the position sensor, and then switched on the high-intensity stimulus trap. This trap was then displaced laterally with respect to the position-sensing trap so that the position sensor reported a displacement of 250 nm. We then compared this nominal displacement to that detected by a camera that acquired brightfield images of the focal plane. The camera had previously been calibrated to accord with well-defined displacements of the nano-positioning stage. Across twelve probe beads, the camera read out an average displacement of 250 ± 13 nm (mean ± SD), values that were in excellent agreement with the microscope’s position sensor. We concluded that the bead-to-bead variability of the calibration was 5 % (Extended Data Table 3c). Because the radius of the probe bead was the parameter with the greatest uncertainty during the calibration procedure, this value also set an upper bound of 5 % on the coefficient of variation of the diameter of our probe beads.

The pedestal bead scattered a small portion of the position sensing beam, which led to an offset of the position signal that was dependent on the pedestal’s position. Although we corrected for this effect through a reference signal, there remained a position offset of ± 4 nm peak-to-peak or 3 nm root-mean-square. This problem contributed uncertainty to the measured displacement between the position-sensing trap and the stimulus trap and thus resulted in reduced accuracy (Extended Data Table 3d). The total accuracy of a nominal distance of 200 nm between the position-sensing trap and stimulus trap was then 10 nm root-mean-square (Extended Data Table 3e). During a single-molecule experiment, the precision of this distance was impacted by the slow relative drift between the two optical traps, which we measured as 3 pm·s^-1^ (Extended Data Table 3f).

Because the spring constant of the stimulus trap depended linearly on the intensity of the stimulus beam, any variation in the beam’s intensity decreased the precision of the spring constant. The spring constant’s root-mean-square noise was 0.27 μN·m^-1^ over 20 s with a vanishingly small drift (Extended Data Table 3g,h). We computed the spring constant from the power-spectral density of the motion of a probe bead as^57^ *k* = 2*πγf_c_*. The error of the drag γ was determined by the uncertainty of the probe bead’s radius (5 %) and exceeded the error of the corner frequency *f_c_*. Depending on the radius of the probe bead, the spring constant of the stimulus trap was therefore accurate to within 5 % of the calibrated value (Extended Data Table 3i).

We determined the force on the trapped probe bead as the product of its extension from the stimulus trap and the trap’s stiffness. Consequently, the precision of the force could be determined from the probe’s position noise, the drift of the stimulus trap, and the noise of the trap’s spring constant. With the probe bead displaced by 200 nm from the stimulus trap and for maximal power of the stimulus laser, we determined a precision of 0.7 pN over 5 min of data acquisition (Extended Data Table 3j). During an experiment the displacement of the probe and the power of the laser were usually smaller than those values, resulting in a smaller uncertainty. The accuracy of the force was determined by the total accuracy of the position of the stimulus trap and that of the spring constant. At maximal power of the stimulus laser and for a displacement of the probe bead from the stimulus trap of 200 nm, we found an experiment-to-experiment uncertainty of the force of 3.8 pN (Extended Data Table 3k). For smaller displacements and lower laser powers this uncertainty was lower.

### Polymer model

We modeled PCDH15 as a freely-jointed chain with *N* = 11 segments, each of length *b*, representing the eleven stiff cadherin domains^34^. The model included in series a worm-like chain^35,59^ characterized by a persistence length *lp*_peptide_ and a contour length *lc*_linkers,total_ = 10 *lc*_linker_, which accounted for the ten flexible disordered linker regions between the stiff domains. Any enthalpic extensibility of the protein was described by a Hookean spring with stiffness *k*_folded_ (Extended Data Fig. 5). Elongation of the protein through the unfolding of each domain was described by the addition of another worm-like chain. Because unfolded polypeptide chains are structurally similar to the inter-domain linker regions, the unfolded portions of the protein were described with an identical persistence length and a contour length of *lc*_unfolded_.

The extension-force relation of the protein was described by the sum of the extension-force relations of the freely-jointed chain and of the worm-like chains,

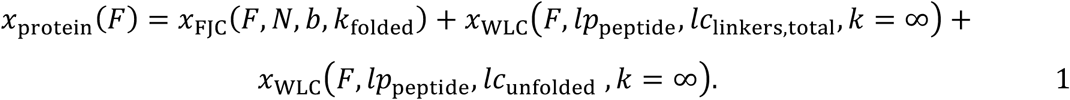

in which the elastic properties of the two worm-like chains were purely entropic (*k* = ∞). The extension-force relation of an extensible freely-jointed chain is given by^34^

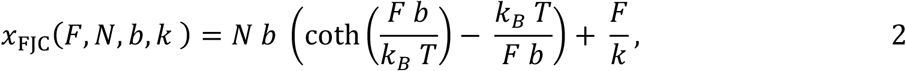

whereas the elongation of an extensible worm-like chain under force is well approximated by^59^

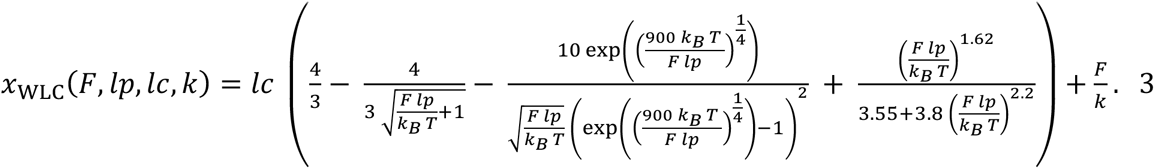

When a force was applied to a single-molecule tether, not only did the tether stretch, but so did the system’s remaining elastic elements, the anchors and linkers in series with the tether (Extended Data Fig. 9). We demonstrated experimentally that the combined mechanics of the assay without PCDH15 could be described by an additional extensible worm-like chain (Extended Data Fig. 10), for which we found across nine experiments an average persistence length *l*p_anchors_ = 0.5 ± 0.1 nm, contour length *l*c_anchors_ = 37 ± 4 nm, and Hookean spring constant *k*_anchors_ = 7.2 ± 1.3 mNm^-1^ (means ± SEMs).

The total polymer model that we fitted to our data was therefore given by

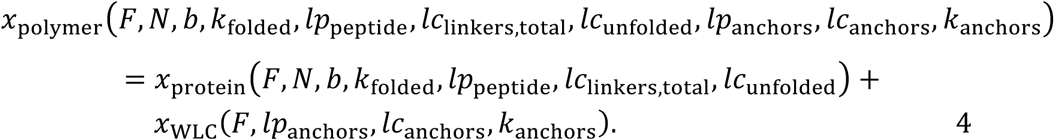

### Determination of the stiffness of PCDH15 and influence of molecular anchors and beads on the measured stiffness

Our force-extension relations capture the mechanics of PCDH15 in series with its molecular anchors and with any compliance of the probe and pedestal beads (Extended Data Fig. 9). To determine the contribution of these elements to our measurements, we conducted experiments without PCDH15 by fusing the amino- and carboxy-terminal anchors and testing their elastic properties in series with the pedestal and probe beads (Extended Data Fig. 10). PCDH15’s stiffness could then be determined by treating the whole system as a series of nonlinear springs,

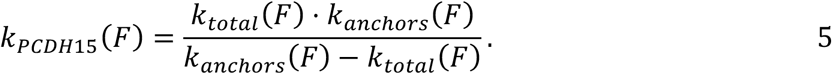

For Ca^2+^ concentrations of 3 mM and 20 μM, we computed the stiffness of the full-length construct—the protein in series with its anchors and the beads—as the spatial derivative of the mean of the force-extension relations associated with its different conformational states (Extended Data Fig. 11). Computed in an analogous way (Extended Data Fig. 10d), the stiffness of the construct without PCDH15 was much larger (Extended Data Fig. 11c,d). From the values of State 1 we computed the stiffness of folded PCDH15 and found that it is surprisingly soft under physiological tensions, offering a stiffness between 0.5 mN·m^-1^ at a tension of 5 pN and 6 mN·m^-1^ at 50 pN (Figure 3a,b).

### Detection of structural changes and fitting of data

The position and force signals (Extended Data Fig 1b,c), sampled at 100 kHz, were split into individual extension-relaxation cycles and smoothed with a second order Savitzky-Golay filter over a window of 101 points, which reduced the temporal resolution to 1 ms. Conformational changes of the protein were automatically detected as sudden increases or decreases in the filtered probe-position signal. To determine whether a structural change occurred at point *i* of The time trace, we computed the averages and standard deviations of 1000 points preceding and succeeding *i*, 〈*x*〉_left_, 〈*x*〉_right_, *σ*_left_, and *σ*_right_. A structural change was detected at point *i* if

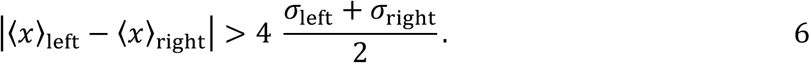

Structural changes that did not fulfill this criterion were missed by our algorithm. If two or more structural changes occurred within 10 ms of one another, our algorithm detected only one larger event.

We then segmented each extension-relaxation cycle into its individual states, demarcated by the structural changes, and fit our polymer model to each of the segments. To facilitate meaningful fits, we sought to constrain the number of free parameters in our polymer model (Equation 4) as much as possible. We first independently measured the parameter values of the worm-like chain describing the compliance of the anchors and of the rest of the assay without PCDH15 (Extended Data Figs. 9 and 10), for which we found an average persistence length *l*p_anchors_ = 0.5 ± 0.1 nm, contour length *l*c_anchors_ = 37 ± 4 nm, and Hookean spring constant *k*_anchors_ = 7.2 ± 1.3 mNm^-1^ (means ± SEMs of nine experiments). We held these values constant at their means for all subsequent fits of extension-relaxation cycles of PCDH15.

In a next step it was important to determine *l*p_peptide_, the persistence length of PCDH15’s unfolded polypeptide chains, for this value entangles the mechanics of State 1 (through the linker regions of the folded protein) with the mechanics of partially unfolded PCDH15. To facilitate this determination, we temporarily approximated State 1 of the protein as a worm-like chain; we then held the parameter values of State 1’s worm-like chain model constant while fitting all extension-relaxation cycles with only *lp*_peptide_ and *lc*_unfolded_ as free parameters, determining *lp*_peptide_ = 0.49 ± 0.04 (mean ± SEM, *N* = 13). Because there was no apparent difference in peptide persistence length between the results for Ca^2+^ concentrations of 3 mM and 20 μM, data were averaged across 13 experiments at both concentrations. This value of *lp*_peptide_ was held constant at its mean for all subsequent fits.

We then fitted State 1 with our protein model to determine the remaining free parameters of the folded protein, *lc*_linker_, *b*, and *k*_folded_ (Table 1). Finally, we fitted all segments of all extension-relaxation cycles with the polymer model, with *lc*_linker_, *b*, and *k*_folded_ held constant at the values determined for the individual proteins, with *lc*_unfolded_ as the only free parameter. The changes of *lc*_unfolded_ between adjacent segments then described the end-to-end elongation of the tether due to unfolding events (Fig. 2).

### Statistical dependence of events A_U_ and B_U_

In order to test whether the order of occurrence of events A_U_ and B_U_ is statistically significant, we simulated extension trials with independent probability densities for the two events such that the resulting force distributions for each of the classes of rips was identical to our experiments. We then showed that the sequence of events in these simulations did not match our experimental observations, proving that the probability densities underlying our experiments are statistically dependent.

For every simulated trial, we first drew two random numbers to determine whether both rips would occur during the extension. The probabilities *p*_A_ and *p*_B_ for respective events A_U_ and B_U_ were determined separately from the experimental data for each PCDH15 molecule. For traces that contained both rips, a parameter representing the pulling force was then linearly increased from 0 pN to 80 pN in steps *dF* = 0.1 pN. After each force increment we allowed rips A_U_ and B_U_ to occur with independent probabilities *ρ*_*A*_(*F*) *dF* and *ρ*_*B*_(*F*)*dF*, in which ρ_A,B_ represent probability densities. Because each class of rips represents a unique structural change, once a rip of either type had occurred no further event of that class could follow. The simulated record was discarded if neither or only one of the rips occurred over the simulated force range. The probability densities *ρ*_*A*,*B*_(*F*) were independently determined for each protein and could be calculated from the experimentally observed force histograms of rips *N*_*A*,*B*_(*F*):

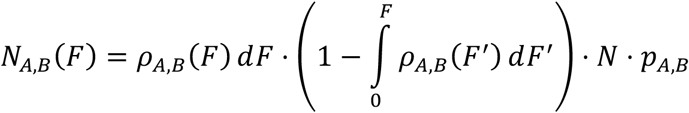

in which *ρ*_*A*,*B*_(*F*) *dF* is the probability of finding rip A_U_ (or B_U_) at force *F* and *N* ⋅ *p*_*A*,*B*_ is the number of extension trials that contain rip A_U_ (or B_U_). The factor in parentheses is the probability that at force *F* during an extension a rip has not yet occurred at forces lower than *F*. This factor accounts for the fact that we draw from *ρ*_*A*,*B*_(*F*) for linearly increasing *F*, and stop drawing from the distribution when a rip occurs. We accordingly can draw from *ρ*_*A*,*B*_(*F*) only if no other rip occurred at a lower force. It follows that

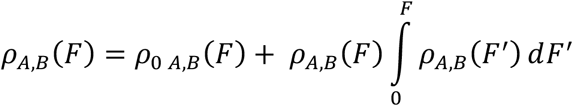

in which 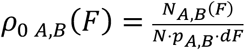 are the normalized force histograms of the experimentally observed rips. We numerically solved this relation to determine *ρ*_*A*,*B*_(*F*) (Extended Data Fig. 12a,c).

Our simulations successfully reproduced the experimentally observed force histograms of the rips A_U_ and B_U_ (Extended Data Fig. 12b,d). We then enumerated the trials in which rip A_U_ occurred before rip B_U_. For the molecule shown in the left column of Figure 2, our simulations— based on the independent probability distributions *ρ*_*A*_(*F*) and *ρ*_*B*_(*F*)—predicted that we should have observed 20.4 extensions in which A_U_ preceded B_U_, a contradiction to the observed value of 8 (*p* < 0.05). Because the counts were assumed to be Poisson distributed, we made statistical comparisons by computing the Wald test statistic 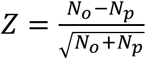, in which *N*_*o*_ are the observed counts and *N*_*p*_ are the predicted counts, followed by a two-tailed comparison of *Z* to a normal distribution to compute the *p*-value. We therefore reject the assumption that A_U_ and B_U_ are independent. An identical conclusion was reached for all five proteins tested at a Ca^2+^ concentration of 3 mM, although two were significant only at *p* < 0.1. When we tested proteins at 20 μM [Ca^2+^] in the same manner, we again routinely observed fewer trials in which rip A_U_ occurred before rip B_U_ than predicted by our simulations. At this physiological Ca^2+^ concentration, however, the difference between the observed and expected number of sequences AB was no longer statistically significant at *p* < 0.1.

### Transformation of force histograms into force-dependent rate constants

We transformed the measured distribution of rip forces of type C_U_ into force-dependent unfolding rate constants^28^. PCDH15 consists of 11 domains that we approximated as equivalent for this analysis. In this approximation, they are each assumed to be equally likely to unfold under force. Hence, a fully folded PCDH15 molecule is eleven times more likely to unfold a domain than a PCDH15 molecule with only one folded domain. To arrive at correct values for the unfolding rates of individual cadherin domains, we therefore computed individual force histograms of C_U_ events originating from each of the states of the protein. We then weighted the measured force histograms with the number of folded cadherin domains in each state: for example, the force histogram for domain unfolding from States 1 and 2, which both correspond to PCDH15 with eleven (at least partially) folded cadherin domains, was divided by *n* = 11 domains. The weighted force histograms were then combined and the remaining analysis performed^28^. Our weighting of the force histograms is similar to the weighting that has been employed in pseudo dwell time analysis for the determination of un- and refolding rates in chains of identical titin domains^33^.

## Extended Data

**Extended Data Fig. 1.**
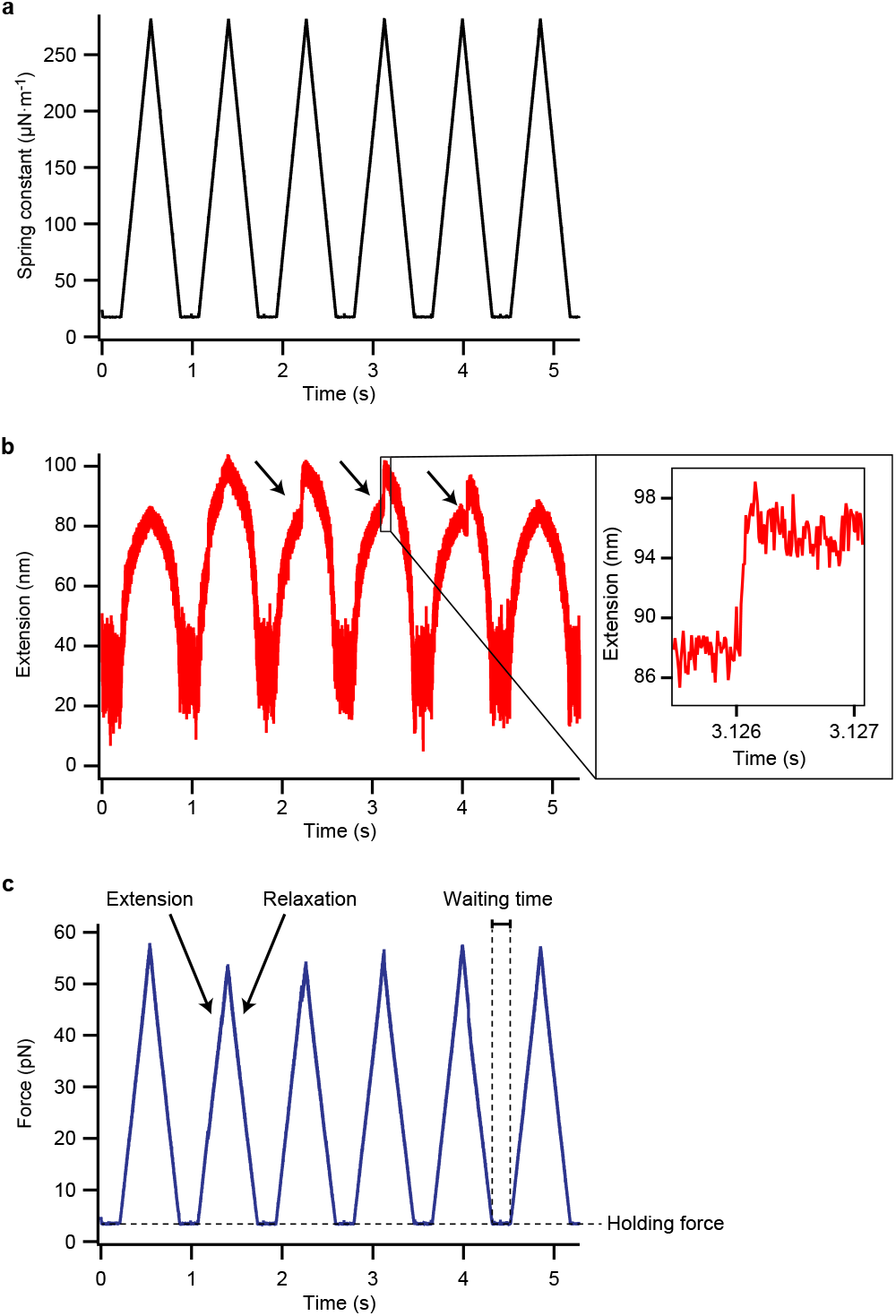
Time traces of extension-relaxation cycles. (**a**) The force on a PCDH15 tether was varied by adjustment of the spring constant of the stimulus trap, which was centered about two hundred nanometers from the equilibrium position of the probe bead. (**b**) The tether’s extension included occasional abrupt events (arrows). Note the extensive noise owing to thermal excitation of the molecule. (**c**) The force acting on the tethered protein was computed from the probe bead’s position and the spring constant of the stimulus trap. Each protein underwent hundreds of extension-relaxation cycles, between which the force was held constant for a particular waiting time so that any unfolded domains could refold. The Ca^2+^ concentration was 3 mM.

**Extended Data Fig. 2.**
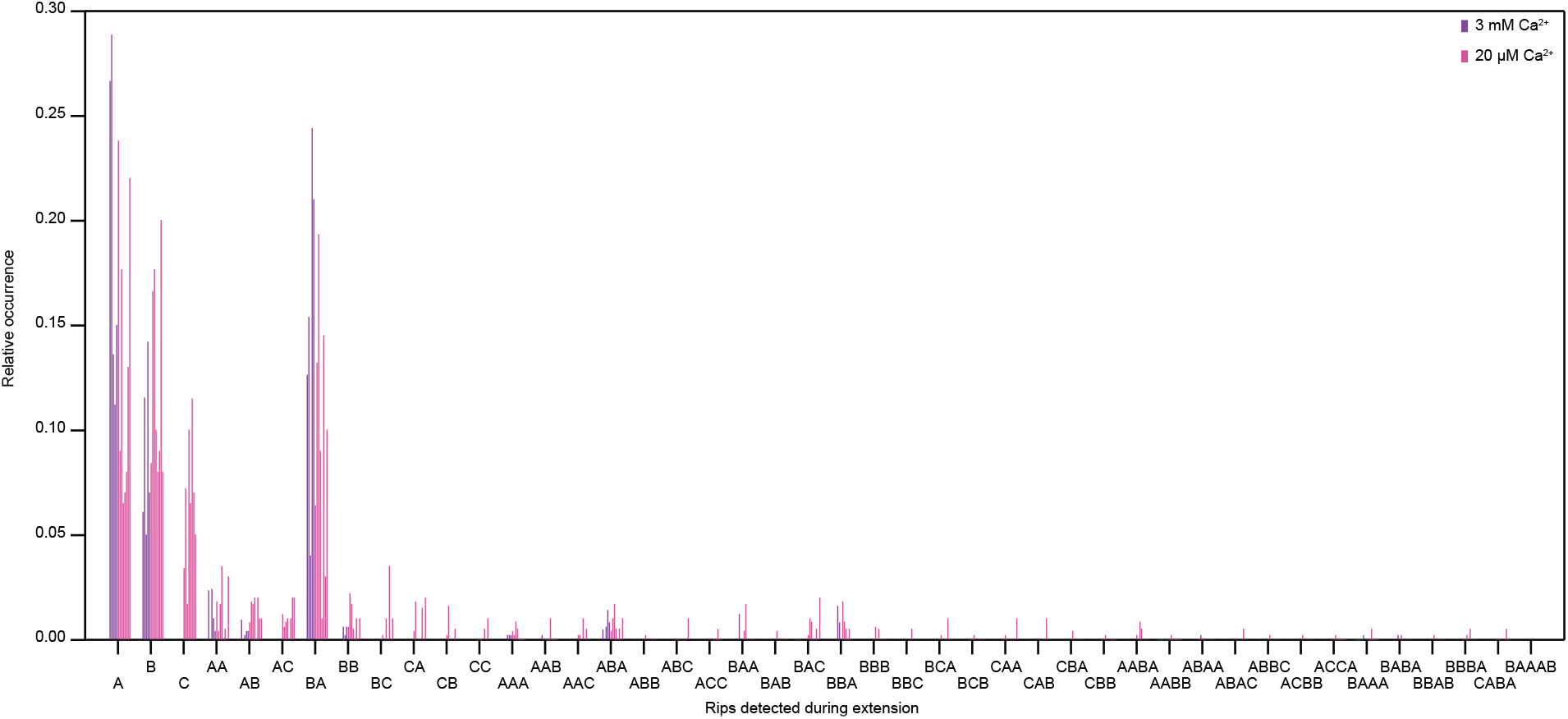
Sequences of structural changes during extension trials. The histogram indicates the fraction of extension traces that contained a specific sequence of unfolding events. For each possible sequence of events, each bar in a cluster represents a single molecule. Each molecule was extended at a loading rate of 130 pN·s^-1^. Depending on the protein, the holding force between cycles was 2-4 pN and the waiting time was 0.2-2 s. This inconsistency altered the extent to which a particular molecule refolded between cycles and added variability to the results.

**Extended Data Fig. 3.**
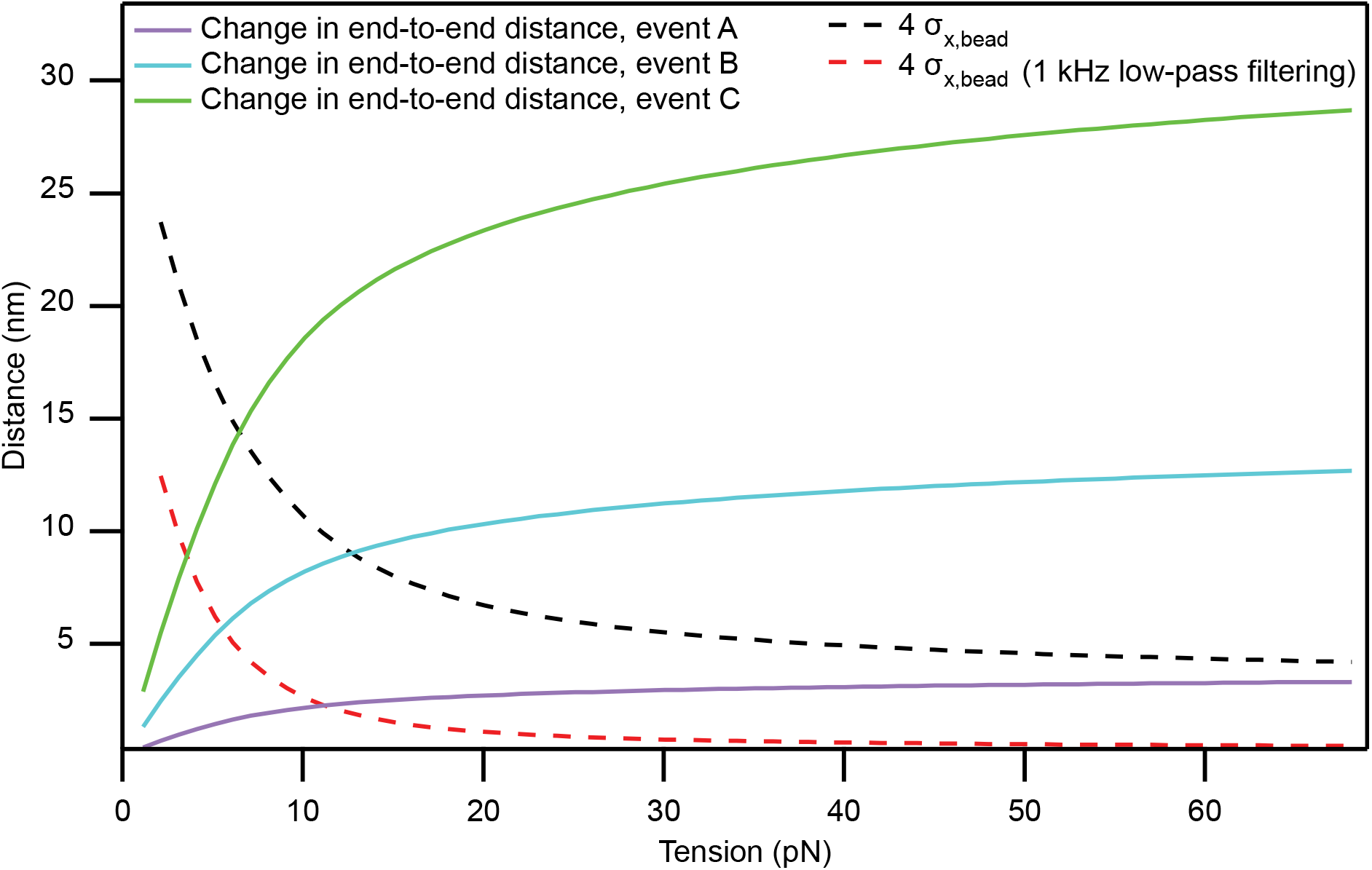
Effect of thermal noise on detectability of conformational changes. When a molecular tether is subjected to a low tension, unfolding or refolding of a domain results in a change in the end-to-end distance much smaller than the actual change in contour length. The expected changes (solid lines) are shown for events of types A, B, and C. If these changes are comparable to the thermal motion of the bead, they cannot be reliably detected by our method. We can confidently identify a folding event if the change in end-to-end distance is larger than four times the standard deviation of the thermal motion (black dashed line, 1 MHz bandwidth; dashed red line, signal low-pass filtered to 1 kHz). The intersection of the dashed and solid lines therefore defines the force below which a given structural change can no longer be reliably resolved for a particular temporal resolution. The band-limited thermal noise in the probe bead’s position was computed as the integral of the power-spectral density of the bead’s motion^57,53^.

**Extended Data Fig. 4.**
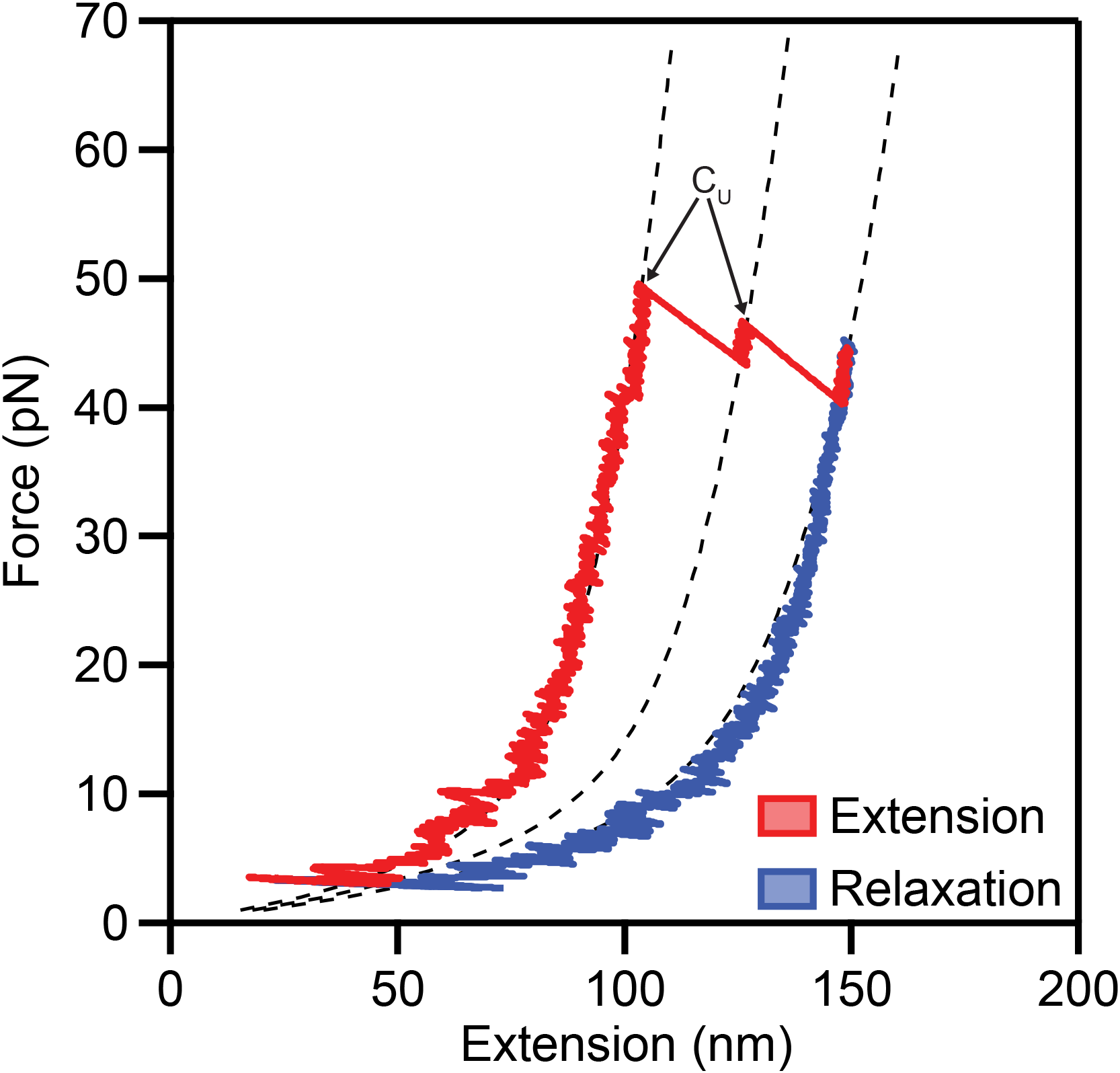
Unfolding of several cadherin domains during one extension-relaxation cycle. In some extension-relaxation cycles of the protein from Fig. 2b, two cadherin domains unfolded during the extension (arrows). The Ca^2+^ concentration was 20 μM.

**Extended Data Fig. 5.**
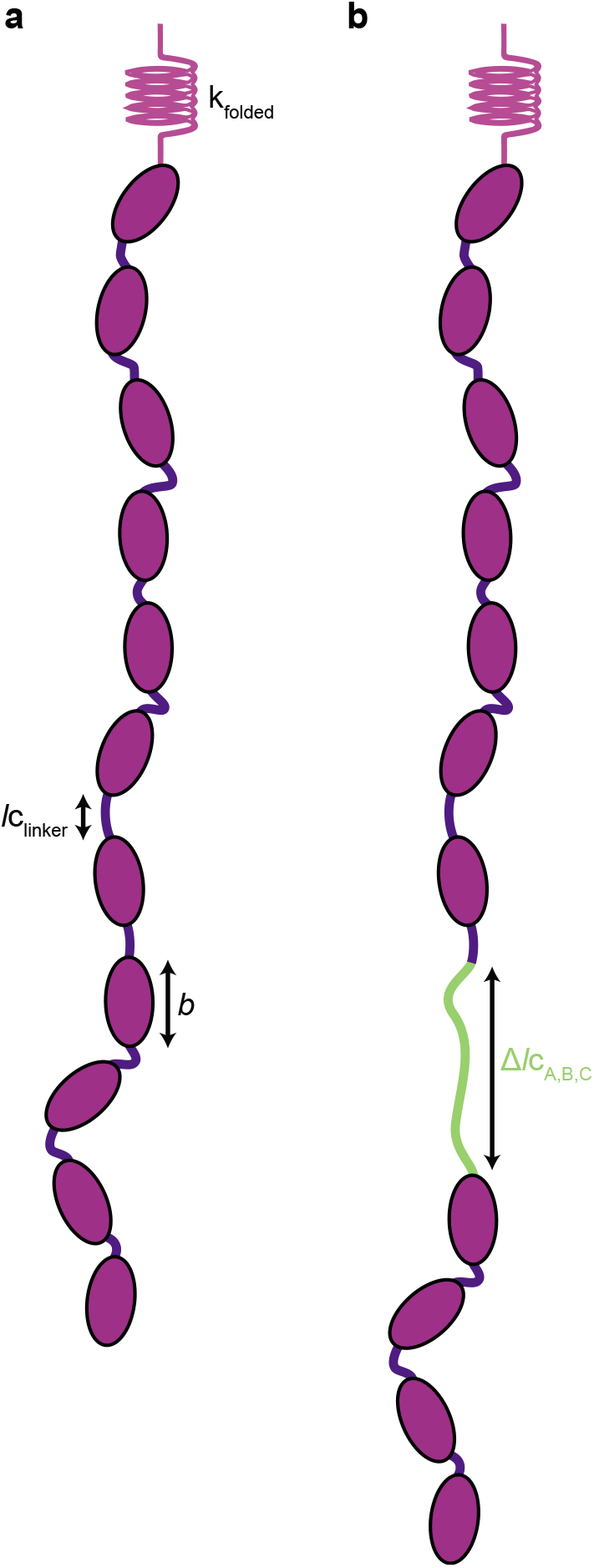
Protein model for PCDH15 under tension. (a) We modeled a folded monomer as a freely jointed chain of eleven stiff segments, each with length *b*. The linker regions between stiff segments consist of unstructured peptides of length *lc*_linker_, whose combined effect was modeled as a worm-like chain with a contour length of 10·*lc*_linker_ and persistence length *l*p = 0.49 nm in series with the freely-jointed chain. A Hookean spring with stiffness *k*_folded_ represents the enthalpic extensibility of the protein. (**b**) Each unfolding event was represented as an additional worm-like chain (green), of the appropriate contour length and of persistence length 0.49 nm, in series with the folded protein.

**Extended Data Fig. 6.**
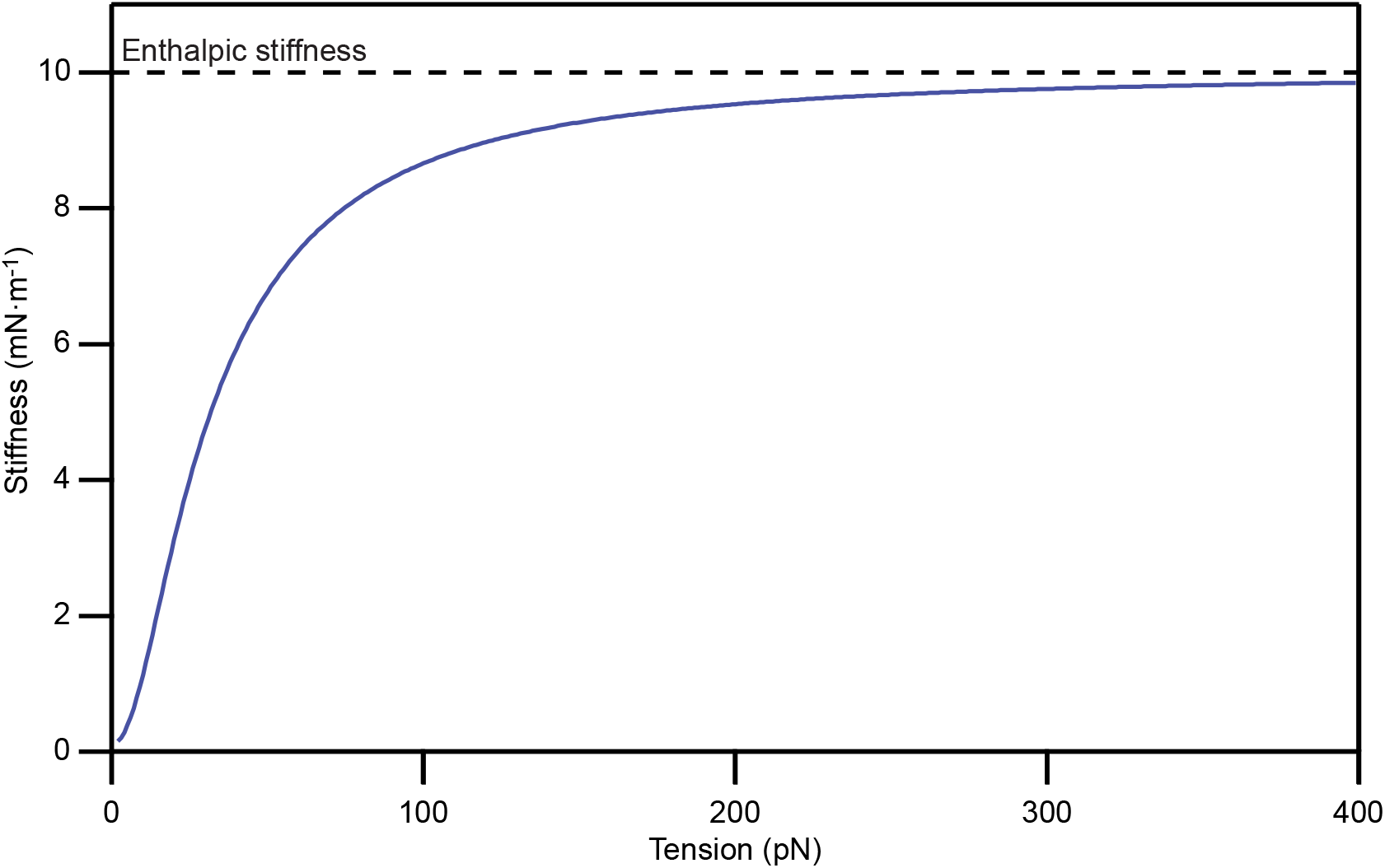
Predicted stiffness of PCDH15 at unphysiologically high forces. Our model predicts that monomeric PCDH15 reaches its enthalpic stiffness only for tensions exceeding hundreds of piconewtons. The physiological range of tensions, 4 pN to 25 pN per molecule, is dominated by the protein’s entropic elasticity.

**Extended Data Fig. 7.**
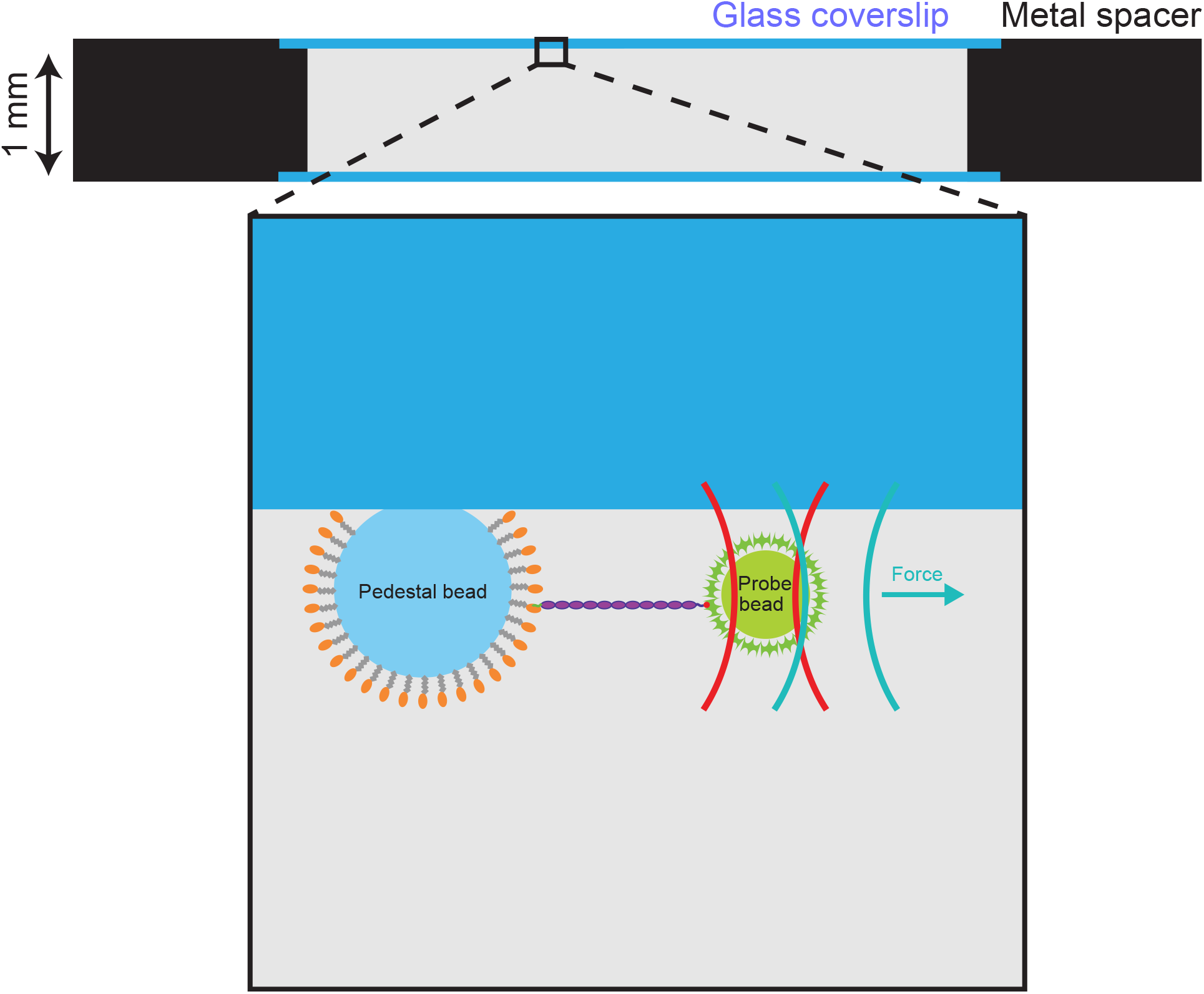
Design of the sample chamber. The sample chamber consisted of two glass coverslips attached by vacuum grease to a metal spacer. The sparsely distributed pedestal beads were covalently attached to the functional surface of the upper coverslip, and the chamber was filled with buffer solution containing freely diffusing probe beads. Note that the photonic-force microscope was of upright design, with the objective lens positioned above the sample chamber.

**Extended Data Fig. 8.**
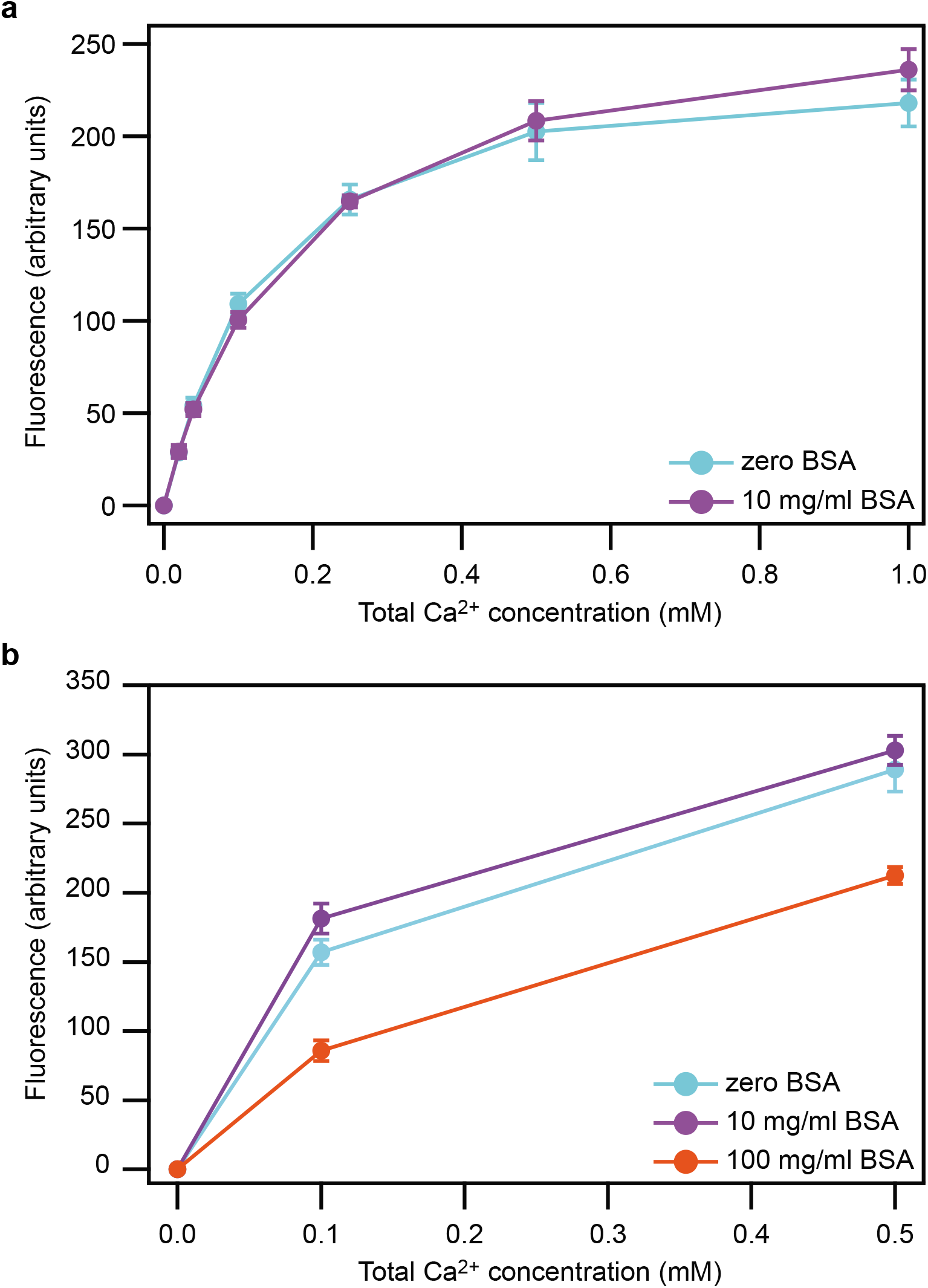
Lack of effect of bovine serum albumin (BSA) on the concentration of free Ca^2+^ ions. (**a**) The fluorescence of the Ca^2+^ indicator Fluo-5N (F14203, ThermoFisher Scientific, Waltham, MA, USA) is unchanged in the presence of 10 mg/ml BSA. (**b**) The concentration of free Ca^2+^ is diminished only by the presence of a BSA concentration as great as 100 mg/ml. Data are plotted as means ± SDs for three experiments.

**Extended Data Fig. 9.**
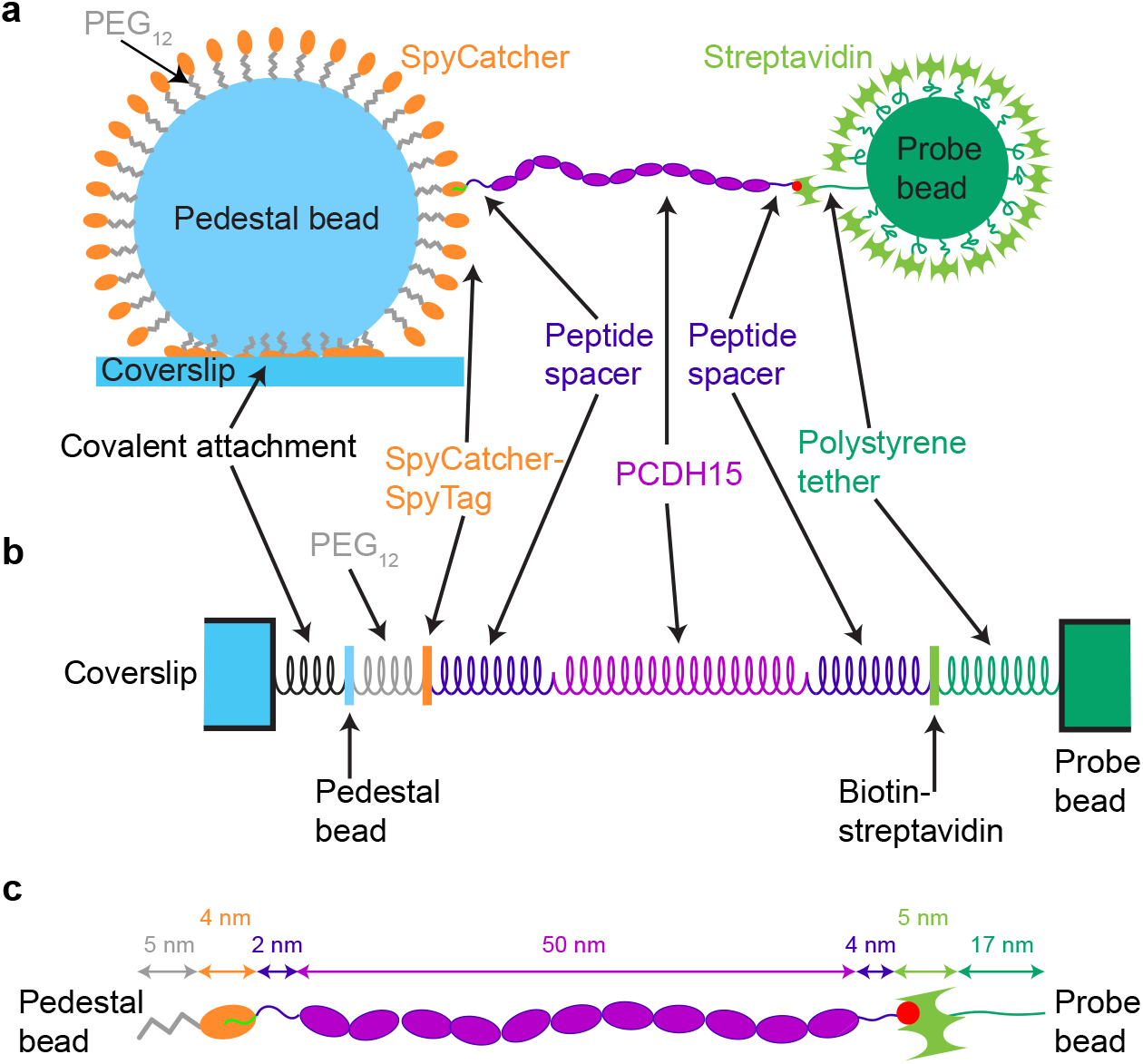
Compliant elements in the single-molecule assay. (**a**) To test the stiffness of PCDH15, we confined an individual monomer between a pedestal and a probe bead. Because each element of the single-molecule assay is compliant, however, the stiffness of the rest of the system—without the PCDH15—must be known in order to accurately determine the protein’s stiffness. The components are not drawn to scale. (**b**) The assay’s compliant elements include the covalent anchoring of the glass pedestal bead to the coverslip, the linkage between the pedestal bead and carboxy-terminus of the protein (polyethylene glycol [PEG], SpyCatcher, SpyTag, and peptide), and the connection between the protein’s amino-terminus and the probe bead (peptide, biotin, and streptavidin). The latter linkage might also include a short polystyrene tether that extends from the probe bead’s surface. (**c**) We designed the assay to contain anchor and linker elements that were as short and stiff as possible. Their approximate contour lengths are indicated. The size of the SpyCatcher, PCDH15, and streptavidin proteins were estimated from their crystal structures (PDB IDs 4MLI, 6CV7, and 1AVD). The lengths of the PEG and peptides are design lengths, and the length of the polystyrene tether was estimated from control experiments (Extended Data Fig. 10).

**Extended Data Fig. 10.**
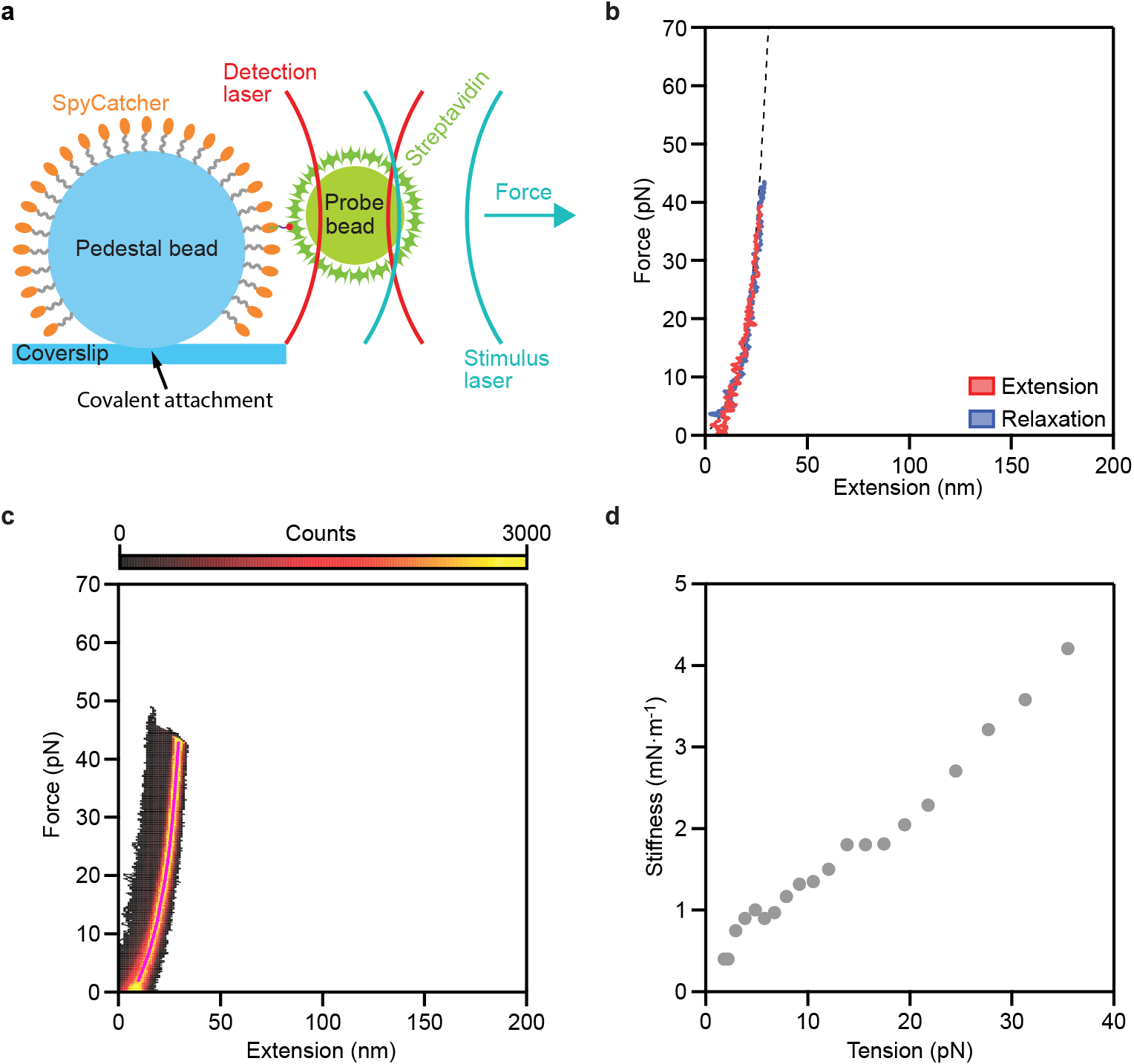
Mechanical properties of the anchors and linkers. (**a**) In order to measure the stiffness of the single-molecule assay system in the absence of PCDH15, we connected the molecular handles (SpyTag and biotinylation peptide) through an eight-amino-acid, flexible linker, anchored this short peptide between a pedestal and a probe bead, and determined its force-extension relation. (**b**) A representative extension-relaxation cycles features a small amount of extensibility that is well fit by a wormlike-chain model. Across nine experiments we found an average persistence length *l*p_anchors_ = 0.5 ± 0.1 nm, contour length *l*c_anchors_ = 37 ± 4 nm, and Hookean spring constant *k*_anchors_ = 7.2 ± 1.3 mNm^-1^ (means ± SEMs). The designed contour length of our tether including PEG, SpyCatcher, peptide, and streptavidin is 20 nm, shorter than the tether’s experimentally determined contour length of 37 nm. We accordingly conclude that the extensible polystyrene hairs on the surface of the probe beads have an average length of 17 nm (Extended Data Fig. 9). (**c**) A state-space heatmap of 500 extension-relaxation cycles shows only the single conformational state expected for an unstructured peptide. These data confirm that neither of our proteinaceous anchors undergoes structural changes over the relevant force range. (**d**) To estimate the compliance of our system of anchors, we determined the average of the highly occupied region in the heat map (pink line in **c**) and computed its slope.

**Extended Data Fig. 11.**
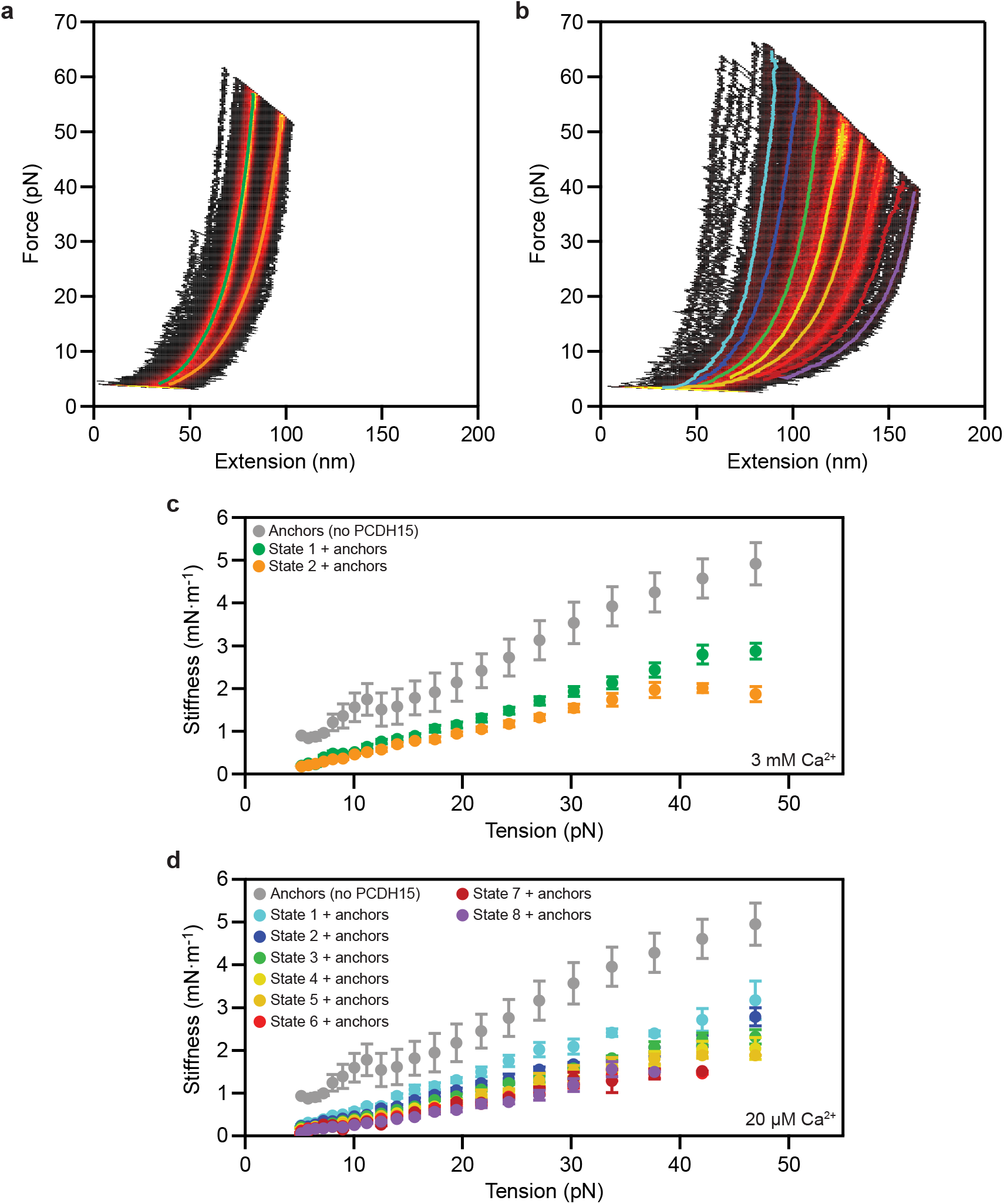
Total stiffness of the single-molecule assay system. (**a**) We determined the stiffness of PCDH15 in its different conformational states at a Ca^2+^ concentration of 3 mM by finding the slope of each highly occupied region of the state-space heatmap. (**b**) The same operation was performed for PCDH15 in the presence of 20 μM Ca^2+^. (**c**) The resulting total stiffnesses represent the compliances of the anchors and of PCDH15 at a Ca^2+^ concentration of 3 mM. (**d**) The result for 20 μM [Ca^2+^] includes six additional unfolded states of progressively diminishing stiffness. As expected, for all states the total stiffness is lower than the stiffness of the anchors alone. By treating PCDH15 and its anchors as springs in series, we can compute from these data the stiffness of PCDH15 alone. The data in **c** and **d** are means ± SEMs for five molecules and six molecules at a Ca^2+^ concentration of respectively 3 mM and 20 μM.

**Extended Data Fig. 12.**
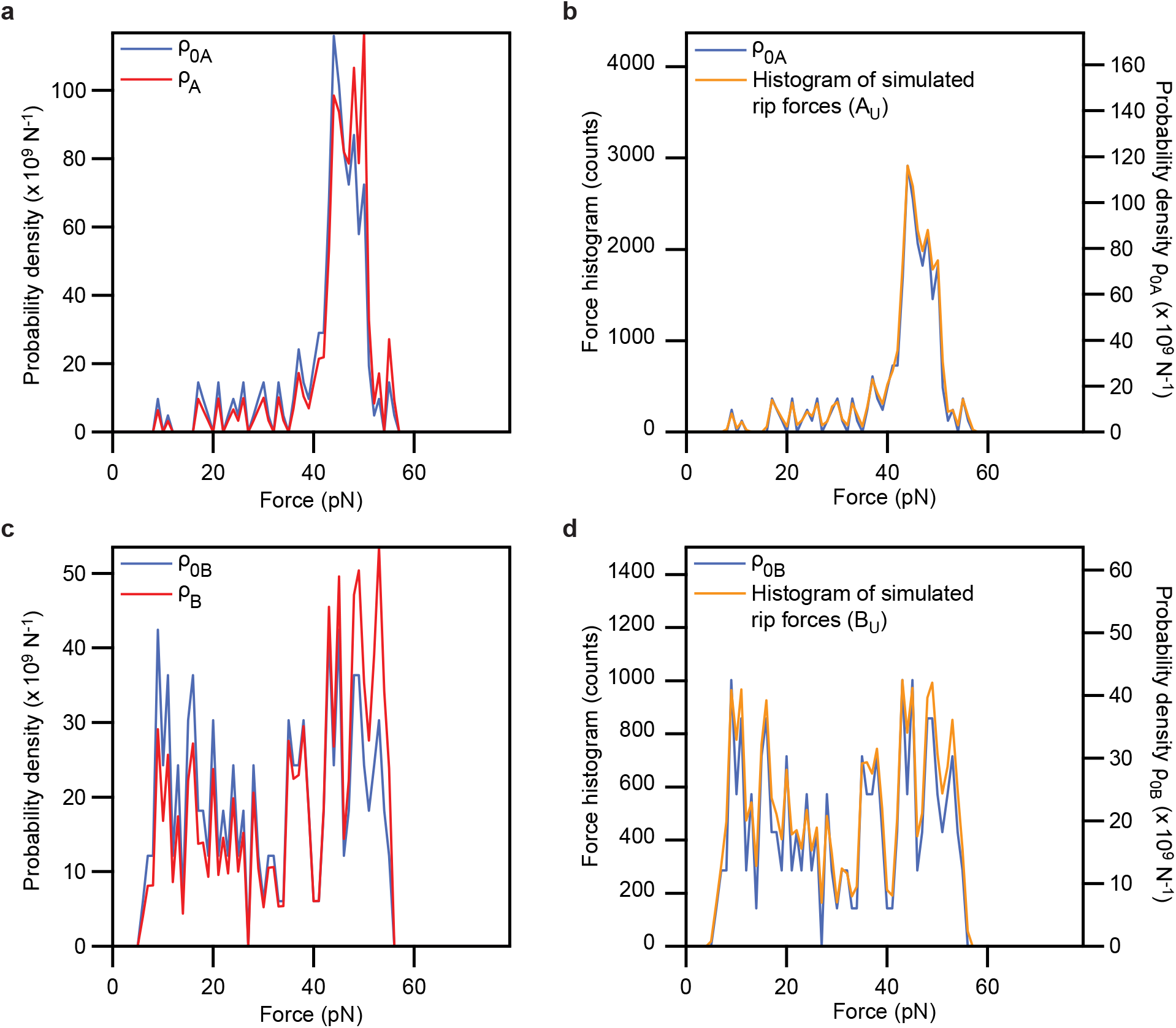
Simulation of the force distributions of the unfolding events A_u_ and B_u_. **(a)** During an extension, a rip of class A occurs with probability *ρ*_*A*_(*F*)*dF*. We calculated the probability density *ρ*_*A*_(*F*) from the experimentally observed force histogram of rips, *ρ*_c*A*_(*F*). **(b)** To confirm that our simulations successfully reproduced the experimentally observed force histograms, we simulated 200,000 extension trials, in each of which rips occurred with a probability *ρ*_*A*_(*F*)*dF*. (**c**) A similar procedure was applied to events of class B. (**d**) A histogram displays the distribution expected from the simulation. For both classes of unfolding event, the histograms of simulated rip forces match the experimentally observed force histograms.

**Extended Data Table 1.**
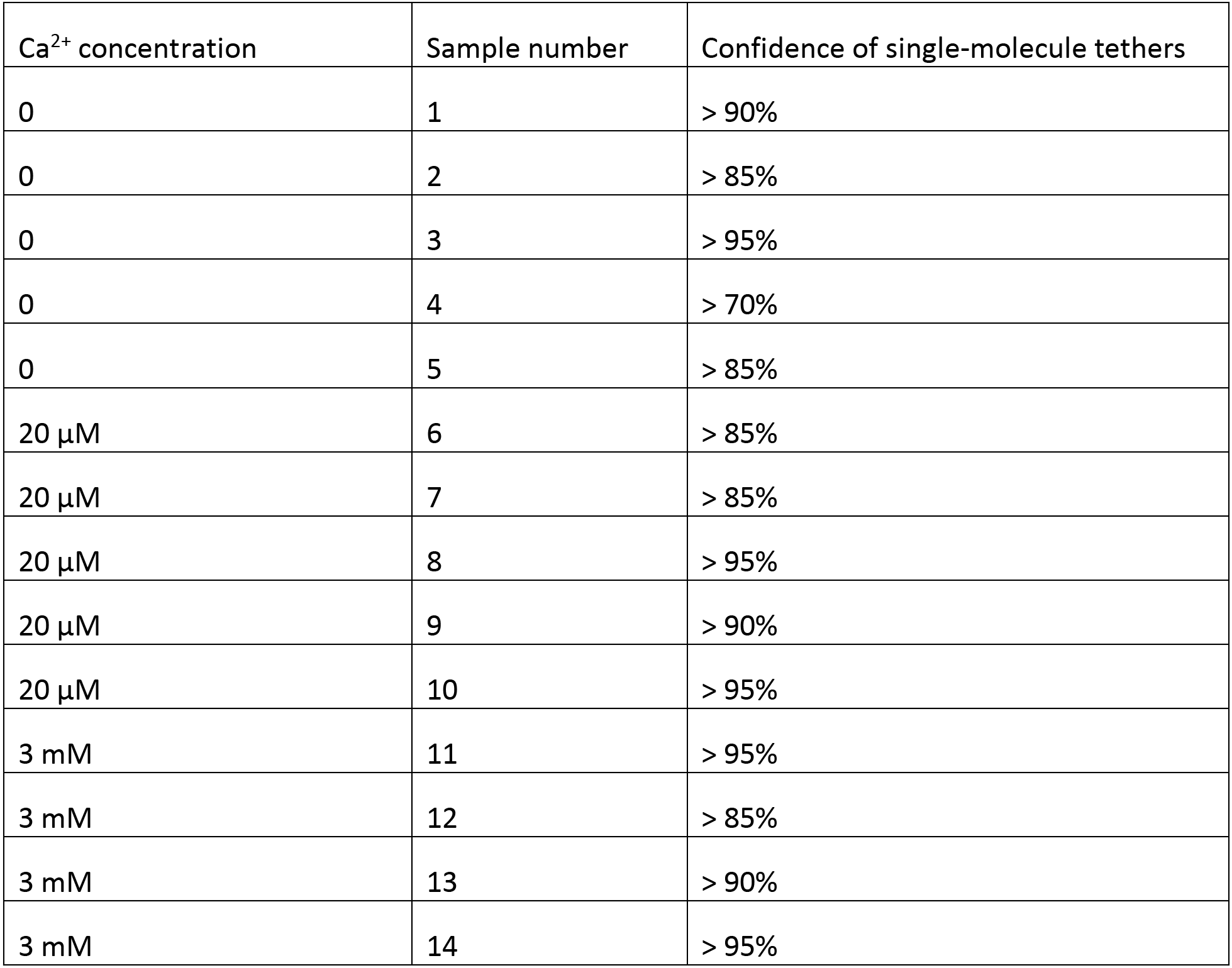
Confidence of tethering a single PCDH15 molecule rather than several^58^.

**Extended Data Table 2.** Confinement of PCDH15 molecules through biotin-streptavidin and SpyCatcher-SpyTag interactions. We tested whether PCDH15 was specifically confined between pedestals and probe beads through its amino- and carboxy-terminal tags. Tether formation was abolished if components of either the SpyCatcher-SpyTag or of the biotin-streptavidin pair were missing.

|  | SpyCatcher-positive<br>Biotinylation-positive |  | SpyCatcher-negative<br>Biotinylation-positive |  | SpyCatcher-positive<br>Biotinylation-negative |  |
| --- | --- | --- | --- | --- | --- | --- |
|  | Field of<br>view 1 | Field of<br>view 2 | Field of<br>view 1 | Field of<br>view 2 | Field of<br>view 1 | Field of<br>view 2 |
| Number of<br>pedestals | 146 | 163 | 89 | 137 | 188 | 168 |
| Number of<br>probe beads | 513 | 493 | 0 | 0 | 0 | 0 |
| Mean probe<br>beads per<br>pedestal | 3.5 | 3.0 | 0 | 0 | 0 | 0 |

**Extended Data Table 3.**
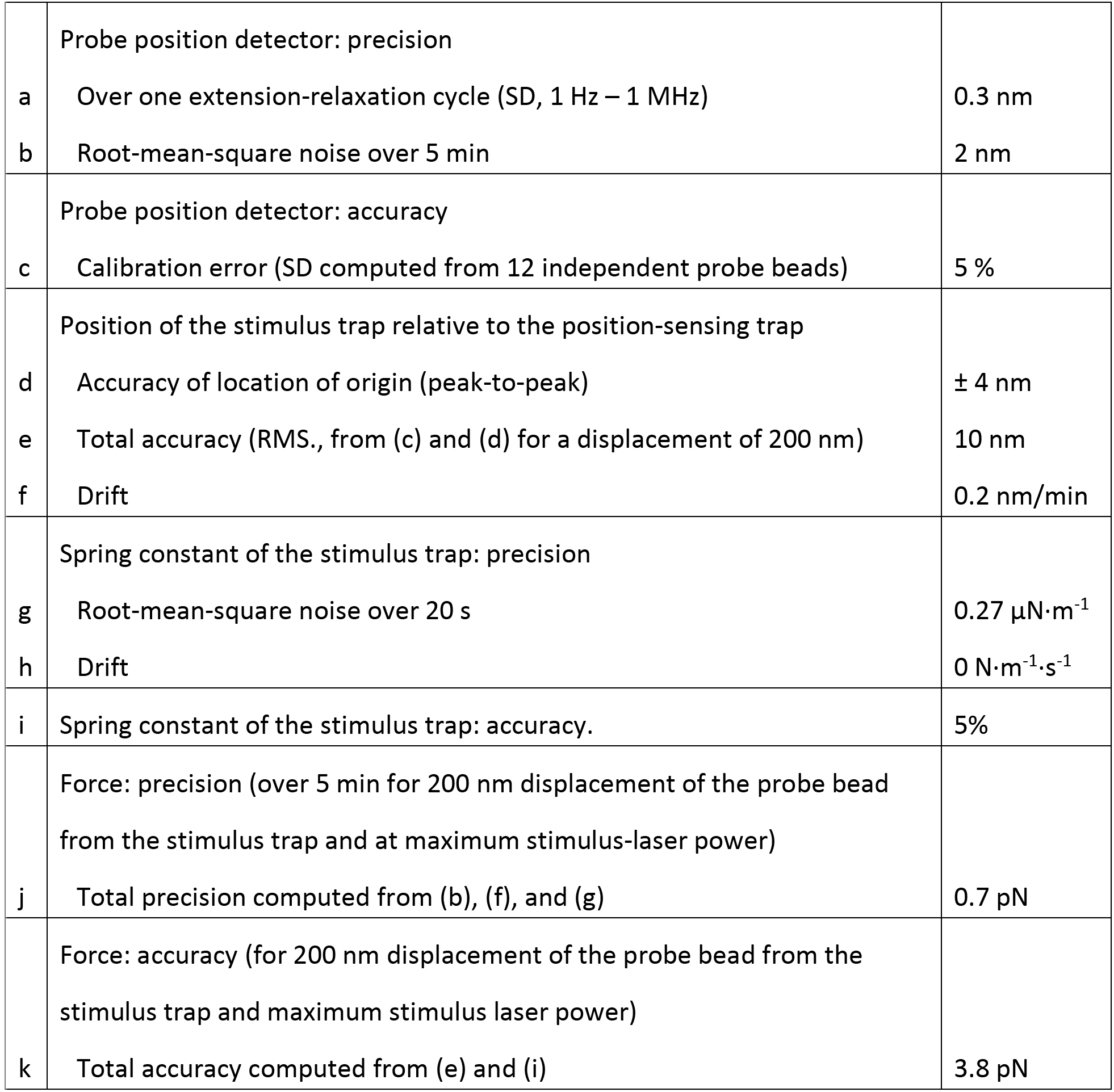
Sources of uncertainty in the photonic-force microscope and single-molecule experiment.

## Extended Data Notes

**Extended Data Note 1 | Interpretation of State 1 as folded PCDH15**

We verified that conformational State 1, that with the smallest contour length (Figure 2d,e), corresponds to fully folded PCDH15. First, we did not observe any reproducibly accessible states with contour lengths shorter than that of State 1. Although we sporadically observe individual force-extension relations to the left of State 1, these curves were not reproducible and likely reflected nonspecific interactions between the pedestal or probe bead surfaces and the protein. Second, fits of our polymer model to the data for each State 1 yielded contour lengths of 46 ± 7 nm and 47 ± 7 nm for Ca^2+^ concentrations of respectively 3 mM and 20 μM (means ± SEMs for respectively five and eight molecules). These values accord well with a contour length of 50 nm expected for a chain of 11 cadherin domains, each 4.5 nm in length^23^.

**Extended Data Note 2 | Influence of possible molten-globule states on domain-unfolding rates.**

At tensions below 20 pN and with Ca^2+^ concentrations of 20 μM or zero, the rate of unfolding of cadherin domains diverges from an exponential relationship and systematically reaches higher values than expected (Figure 4). What causes this effect? It is possible that cadherin domains refold through a long-lived molten-globule state, as occurs for other proteins with immunoglobin-like folds^60^. During the waiting time between trials, an unfolded domain might refold completely, fold into the molten-globule state, or not refold at all. If folding occurred, the domain might unfold during the next extension from the completely folded state or from the molten-globule state. We cannot differentiate between these two possibilities, for they would result in indistinguishable contour-length changes. Molten globules are known to unfold readily at low forces^60,61^. Because such rapid unfolding events would increase our measured unfolding rates, they might explain the unexpected behavior observed at low tensions.

**Extended Data Note 3 | An estimate for the enthalpic stiffness of full-length, dimeric tip links.**

A full-length tip link consists of a dimer of PCDH15, which forms the lower one-third of the filament, and a dimer of CDH23 that makes up the upper two-thirds of its length. For monomeric PCDH15 we found an enthalpic stiffness of 9 mN m^-1^. A PCDH15 dimer, made up of two monomers in parallel, then has a stiffness of roughly 18 mN m^-1^. If CDH23 has a similar enthalpic stiffness per length, then the full-length dimeric tip link has a stiffness of one-third of that of dimeric PCDH15, resulting in an estimate of 6 mN m^-1^.

